# Bone marrow adipogenic lineage precursors (MALPs) promote osteoclastogenesis in bone remodeling and pathologic bone loss

**DOI:** 10.1101/2020.08.01.231829

**Authors:** Wei Yu, Leilei Zhong, Lutian Yao, Yulong Wei, Tao Gui, Ziqing Li, Hyunsoo Kim, Nathaniel Dyment, Xiaowei S. Liu, Shuying Yang, Yongwon Choi, Jaimo Ahn, Ling Qin

**Affiliations:** Department of Orthopaedic Surgery, Perelman School of Medicine, University of Pennsylvania, Philadelphia, PA 19104, USA; Department of Orthopaedics, Union Hospital, Tongji Medical College, Huazhong University of Science and Technology, Wuhan 430022, China; Department of Bone and Joint Surgery, Institute of Orthopedic Diseases, The First Affiliated Hospital, Jinan University, Guangzhou, Guangdong, China; Department of Basic & Translational Sciences, School of Dental Medicine, University of Pennsylvania, Philadelphia, PA 19104, USA; Department of Pathology and Laboratory Medicine, Perelman School of Medicine, University of Pennsylvania, Philadelphia, PA 19104, USA

## Abstract

Bone is maintained by coupled activities of bone-forming osteoblasts/osteocytes and bone-resorbing osteoclasts and an alternation of this relationship can lead to pathologic bone loss such as in osteoporosis. It is well known that osteogenic cells support osteoclastogenesis via synthesizing RANKL. Interestingly, our recently identified bone marrow mesenchymal cell population—marrow adipogenic lineage precursors (MALPs) that form a multi-dimensional cell network in bone—was computationally demonstrated to be the most interactive with monocyte-macrophage lineage cells through highly and specifically expressing several osteoclast regulatory factors, including RANKL. Using an adipocyte-specific *Adipoq-Cre* to label MALPs, we demonstrated that mice with RANKL deficiency in MALPs have a drastic increase of trabecular bone mass in long bones and vertebrae starting from 1 month of age but that their cortical bone is normal. This phenotype was accompanied by diminished osteoclast number and attenuated bone formation at the trabecular bone surface. Reduced RANKL signaling in calvarial MALPs also abolished osteolytic lesions after lipopolysaccharide (LPS) injections. Furthermore, in ovariectomized mice, elevated bone resorption was partially attenuated by RANKL deficiency in MALPs. In summary, our studies identified MALPs as a critical player in controlling bone remodeling during normal bone metabolism and pathological bone loss in a RANKL-dependent fashion.

## Introduction

Bone is a dynamic tissue, constantly undergoing remodeling through coupled activities of bone-resorbing osteoclast and the bone-forming osteoblast/osteocyte (1). A shift in the balance of these two actions toward resorption leads to osteoporosis, an insidious disease characterized by excessive bone loss and micro-architectural deterioration leading to fragility and increased risk of fracture (2). As a highly prevalent disorder, osteoporosis affects more than 75 million people in the US, Europe and Japan and is the underlying condition related to more than 8.9 million fractures annually worldwide (3).

Mature osteoclasts are large multinucleated cells derived from the monocyte-macrophage lineage of hematopoietic origin (4). They firmly attach to the bone surface and degrade bone matrix. Osteoclast differentiation predominantly depends on the signal from RANKL (encoded by *Tnfsf11* gene), a type II transmembrane protein of the TNF superfamily, and is modulated by other cytokines and growth factors (5). *Tnfsf11*^−/−^ mice have no osteoclasts in bone and exhibit severe osteopetrosis (high bone mass) phenotype (6, 7). The early studies indicated that osteoblasts and their progenitors are the major source of RANKL in bone to support osteoclastogenesis (8). Later, animal studies showed that osteoblast ablation does not affect osteoclast formation (9, 10). In growing mice, hypertrophic chondrocytes appear to be the main source of RANKL for bone resorption (11). In adult mice, osteocytes, the descendants of osteoblasts that are embedded in the bone matrix, have been demonstrated to be the major stimulator of osteoclastogenesis (11-13).

Osteoblasts and osteocytes are derived from bone marrow mesenchymal stem cells (MSCs), which also give rise to marrow adipocytes. Recently, we computationally delineated the hierarchy of mesenchymal lineage cells from MSCs to mature cells using large scale single cell RNA-sequencing (scRNA-seq). Surprisingly, this study unveiled a new cell population, marrow adipogenic lineage precursors (MALPs), situating along the adipogenic differentiation route after mesenchymal progenitors and before classic lipid-laden adipocytes (LiLAs) (14). Labeled by mature adipocyte-specific *Adipoq-Cre* (15, 16), MALPs are abundant, non-proliferative cells that express many adipocyte markers but have no lipid accumulation. Shaped as a central body with multiple cell processes, they exist as stromal cells and pericytes forming a 3D network ubiquitously distributed within the bone marrow and function to maintain vessel structure and to inhibit bone formation.

Our scRNA-seq datasets were derived from analyzing Td^+^ cells sorted from endosteal bone marrow of *Col2-Cre Rosa-tdTomato* (*Col2/Td*) mice at various ages. In these mice, Td labels the entire mesenchymal lineage cells (14, 17, 18). Interestingly, our datasets also contained many hematopoietic cells, including osteoclasts. While these “contaminant” cells did not affect the single cell analysis of mesenchymal lineage cells, they actually provided an advantage for delineating the osteoclast differentiation process and for examining the interaction between osteoclastogenesis and mesenchymal subpopulations. To our surprise, our in silico data indicated that MALPs, not osteoblasts or osteocytes, are the most supportive cells for osteoclast formation. To validate this finding, we constructed adipocyte-specific *Tnfsf11 CKO* mice to investigate the role of MALP-derived RANKL in regulating bone remodeling at various skeletal sites under physiological and pathological conditions.

## Methods

See supplementary methods for details.

## Results

### ScRNA-seq analysis reveals osteoclastogenesis in bone marrow

To identify bone marrow mesenchymal subpopulations and to delineate the bi-lineage differentiation paths of bone marrow MSCs, we performed scRNA-seq on top 1% Td^+^ cells sorted from endosteal bone marrow of 1-3-month-old *Col2/Td* mice. Combining three batches of sequencing data together generated a dataset containing 17494 cells with 2519 genes/cell and 11071 UMIs/cell. Clustering analysis revealed 20 cell clusters (Fig. 1A), including 8 mesenchymal lineage cell clusters (Fig. S1A), 10 hematopoietic lineage cell clusters, 1 endothelial cell cluster, and 1 mural cell cluster. Our previous study annotated mesenchymal clusters as early mesenchymal progenitors (EMPs), intermediate mesenchymal progenitors (IMPs), late mesenchymal progenitors (LMPs), lineage committed progenitors (LCPs), osteoblasts, osteocytes, MALPs, and chondrocytes (14). Based on known lineage markers, hematopoietic cells were divided into hematopoietic stem and progenitor cells (HSPCs), monocyte progenitors (MP), granulocyte progenitors (GP), granulocytes, B cells, T cells, erythrocytes (Fig. S1B), monocytes, macrophages, and osteoclasts (Fig. 1B). Hierarchy analysis showed distinct gene expression signature in each cluster (Fig. S1C).

**Figure 1.**
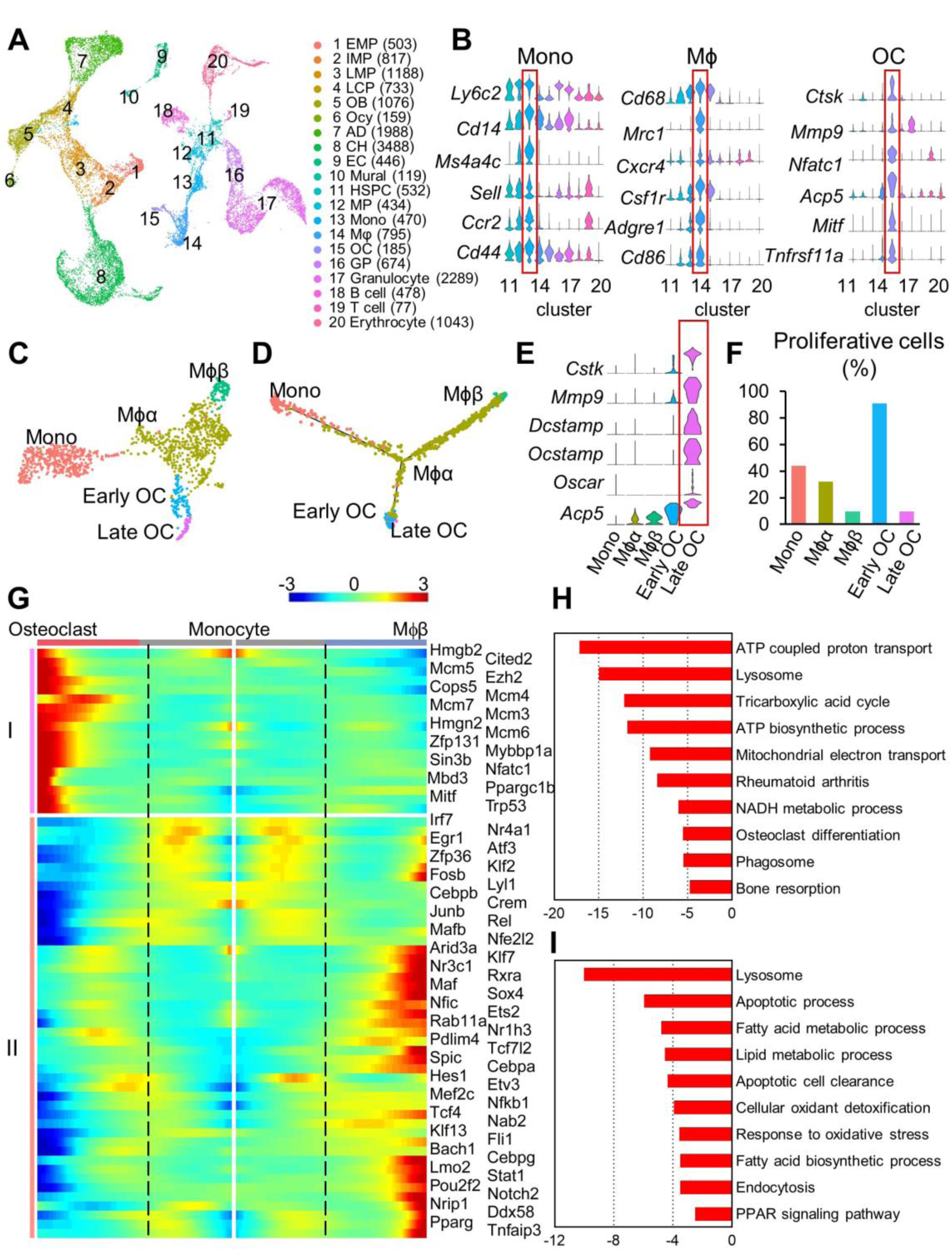
Single cell RNA sequencing identifies bone marrow monocyte-macrophage lineage cells and delineates in vivo osteoclastogenesis. (A) The UMAP plot of cells isolated from bone marrow of 1-3-month-old *Col2/Td* mice (n=5 mice). EMP: early mesenchymal progenitor; LMP: late mesenchymal progenitor; OB: osteoblast; Ocy: osteocyte; LCP: lineage committed progenitor; CH: chondrocyte; EC: endothelial cells; HSPC: hematopoietic stem and progenitor cells; MP: monocyte progenitor; Mϕ: macrophage; OC: osteoclast; GP: granulocyte progenitor. (B) Violin plots of marker gene expression for monocyte, macrophage, and osteoclast clusters. (C) The UMAP plot of monocyte-macrophage lineage cells only. (D) Monocle trajectory plot of monocyte-macrophage lineage cells. (E) The expression patterns of known late osteoclast markers in violin plots. (F) The percentage of proliferative cells (S/G2/M phase) among each cluster was quantified. (G) Pseudotemporal depiction of differentially expressed TFs starting from the branch point (dashed lines) toward osteoclast (left) and macrophage (right) differentiation. Group I and II contain TFs that are highly up-regulated during osteoclast and macrophage differentiation routes, respectively. Color bar indicates the gene expression level. (H) GO term and KEGG pathway analyses of genes up-regulated in late osteoclasts compared to early osteoclasts. (I) GO term and KEGG pathway analyses of genes up-regulated in Mϕβ cells compared to Mϕα cells.

Osteoclasts in postnatal mice are mostly derived from monocyte-macrophage lineage of HSPC descendants. Indeed, monocyte, macrophage, and osteoclast cells (cluster 13, 14, and 15, respectively) were close to each other in the UMAP plot. Separately analyzing these cells using UMAP or Monocle generated 1 monocyte cluster at one end of the pseudotime trajectory, 1 macrophage cluster (Mϕα) at the branch point, 1 macrophage cluster (Mϕβ) at the 2^nd^ end, 1 early osteoclast cluster and 1 late osteoclast cluster at the 3^rd^ end (Fig. 1C, D), suggesting that monocytes undergo bi-lineage differentiation into mature Mϕβ cells and osteoclasts via Mϕα as the intermediate cell type. Genes related to mature osteoclast functions, such as fusion, matrix digestion, and proton translocation, were highly expressed in the late osteoclast cluster (Fig. 1E). Cell cycle analysis confirmed that terminal differentiated Mϕβ cells and late osteoclasts are non-proliferative while other cells, particularly early osteoclasts, are highly proliferative (Fig. 1F).

Positioning individual cells along a linear pseudo-timeline with monocytes as the root revealed known and novel transcription factors (TFs) differentially expressed after the branch point into 2 lineages (Fig. 1G). Nfatc1, a master regulator of osteoclast differentiation (19), was present within the osteoclast lineage differentiation, supporting the reliability of our computational analysis. Other known TFs, such as Ppargc1b (20), Mitf (21), and Ezh2 (22), and Hmgb2 (23), were also identified in this assay. To date, TFs driving macrophage differentiation inside the bone marrow are largely unknown. Our analysis suggested that some adipogenic TFs, such as Pparg, Cebpa, Cebpb etc, and Notch signaling TFs, such as Notch 2 and Hes1, are up-regulated during the differentiation route of macrophages.

GO term and KEGG pathway analyses of differentially expressed genes (DEGs) between early and late osteoclasts pointed out many known features of osteoclasts, such as proton transport, ion transport, ATP biogenesis, mitochondrial related metabolic pathways (Fig. 1H, Table S1). Early osteoclasts were enriched with cell cycle genes, indicating their proliferative nature (Fig. S2A). Comparison of two macrophage clusters indicated that the main function of Mϕβ is efferocytosis because pathways, such as apoptotic cell clearance, endocytosis, oxidation-reduction processes, lipid metabolic etc., were enriched in this cluster of cells (Fig. 1I, Table S2). In contrast, Mϕα was associated with translation, immune-response, and chemotaxis (Fig. S2B), suggesting that these cells are more involved in regulating their environment. Collectively, our scRNA-seq dataset provided a powerful tool for analyzing the in vivo differentiation route of osteoclasts as well as cellular functional changes.

### MALPs are specifically labeled by Adipoq-Cre in adult mice

Our previous study used mature adipocyte-specific *Adipoq-Cre* to label MALPs in 1-monthold mice. Since marker gene expression could be fluid among mesenchymal subpopulations during aging (14), we first investigated whether the same specificity holds in adult mice. For this purpose, we constructed *Adipoq-Cre Tomato* (*Adipoq/Td*) mice with or without *2*.*3kbCol1a1-GFP* (*Col1-GFP*) that labels osteoblasts (24). At 3 months of age, *Adipoq/Td/Col1-GFP* mice showed prominent Td signal inside the bone marrow of long bone (Fig. 2Aa). However, chondrocytes in articular and growth plate cartilage were not labeled (Fig. 2Ab, c). Td^+^ cells were present at the chondro-osseous junction (COJ) between growth plate and primary spongiosa and throughout the entire metaphyseal and diaphyseal bone marrow. Only 4% of GFP^+^ osteoblasts and 1% of GFP^+^ osteocytes were Td^+^ (Fig. 2Ad, e, B), indicating that *Adipoq-Cre* rarely labels osteoblasts and osteocytes in adult mice. We did observe some Td^+^ cells at the bone surface but they were not GFP^+^ (Fig. 2Ad). Furthermore, we did not find any Td^+^ cells at the periosteal surface of cortical bone (Fig. 2Af). Inside the bone marrow, Td^+^ cells existed as pericytes surrounding capillaries (Fig. 2C) and non-hematopoietic stromal cells (Fig. 2D). They did not incorporate EdU (Fig. 2E), suggesting a non-proliferative nature. As expected, all Perilipin^+^ bone marrow adipocytes were Td^+^ (Fig. 2F).

**Figure 2.**
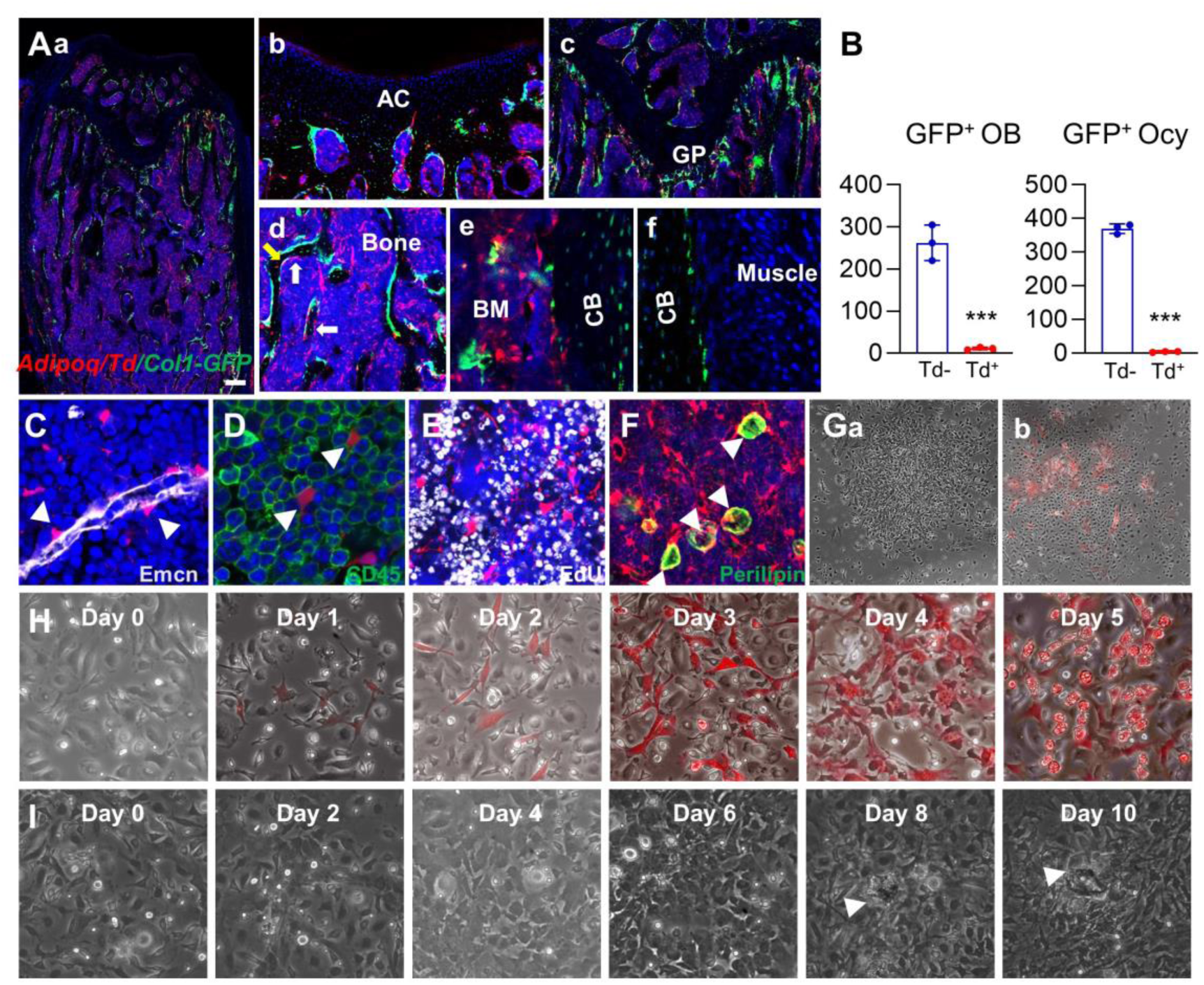
*Adipoq-Cre* labels MALPs in adult mouse bone marrow. (A) Representative fluorescent images of 3-month-old *Adipoq/Td/Col1-GFP* mouse bone reveal many bone marrow Td^+^ cells. (a) A low magnification image of distal femur. Scale bar=200 μm. (b-e) At a high magnification, Td does not label chondrocytes in articular cartilage (b) and growth plate (c), osteoblasts, nor osteocytes (d, e, f). White and yellow arrows point to Td^+^GFP^−^ cells and Td^+^GFP^+^ cells at the bone surface, respectively. (B) Quantification of Td^+^ and Td^−^ cells among GFP^+^ osteoblasts and osteocytes. ***: p<0.001, n=3-5 mice/group. (C) Td labels pericytes (arrow heads) in bone marrow. Emcn: endomucin for vessel staining. (D) In *Adipoq/Td* mice, Td does not label CD45^+^ hematopoietic cells. (E) In vivo EdU injection reveals that bone marrow Td^+^ cells do not proliferate. (F) All Perilipin+ adipocytes (arrow heads) are Td^+^ as well. (G) CFU-F assay of bone marrow cells from *Adipoq/Td* mice shows that all CFU-F colonies are made of Td^−^ cells (a). Some Td^+^ cells do attach to the dish and form a small cluster within a Td^−^ CFU-F colony (b). (H) In vitro adipogenic differentiation of Td^−^ mesenchymal progenitors reveals that Td signal is turned on first followed by lipid accumulation. The same area was imaged daily by inverted fluorescence microcopy. (I) In vitro osteogenic differentiation of Td^−^ mesenchymal progenitors reveals that Td signal remains off during differentiation. The same area was imaged daily by inverted fluorescence microcopy. Arrow heads point to a bone nodule.

To further examine whether *Adipoq-Cre* labels progenitors, we cultured bone marrow mesenchymal progenitors for osteogenic and adipogenic differentiation. As shown in Fig. 2Ga, most CFU-F colonies were made of 100% Td^−^ cells. A few colonies contained some Td^+^ cells but the majority of cells inside the colony were Td^−^ (Fig. 2Gb), indicating that Td^+^ cells lack colony-formation ability and therefore are not proliferative progenitors. When subjected to adipogenic differentiation, Td^−^ progenitors became Td^+^ cells first (day 1-2) and then accumulate lipid droplets (day 4-5) (Fig. 2H). On the contrary, Td^−^ progenitors started to form bony nodules around day 8-10 and maintained as Td^−^ cells during osteogenic differentiation process (Fig. 2I). Since Perilipin^−^Td^+^ cells (MALPs) were 180 times more than Perilipin^+^Td^+^ cells (lipid-laden adipocytes, LiLAs) in bone marrow at this age (3600 MALPs vs 20 MERAs out of 3620 Td^+^ cells counted, n=3 mice), our data demonstrate that *Adipoq-Cre* is suitable to specifically target MALPs, a non-proliferative, committed adipogenic precursor population in bone marrow of adult mice.

### MALPs are the major source of osteoclast regulatory factors

It is well accepted that mesenchymal lineage cells promote osteoclast precursors to differentiate into mature osteoclasts. With the identification of mesenchymal subpopulations in bone, we next sough to find out which of them mostly communicates with monocyte-macrophage lineage cells. To do so, we calculated the number of ligand-receptor pairs between each mesenchymal subpopulation and monocytic subpopulation in our scRNA-seq dataset. Interestingly, MALPs displayed the most interactions with all three monocytic subpopulations, followed by EMPs (Fig. 3A). Surprisingly, osteoblasts and osteocytes had the least number of interactions. Within monocytic cells, macrophages had the most interactions with MALPs, followed by osteoclasts.

**Figure 3.**
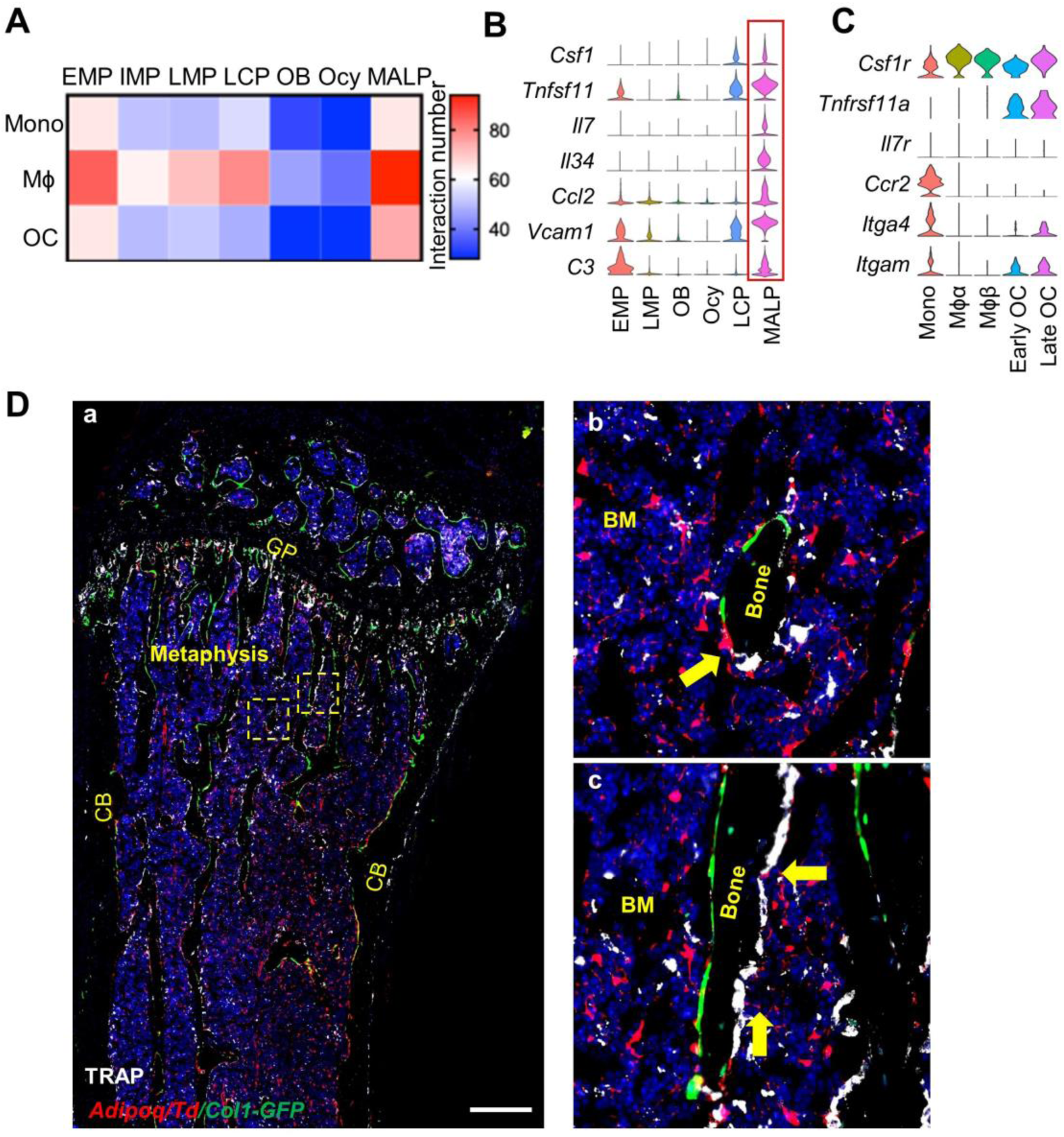
MALPs are the major producer of osteoclast regulatory factors in bone. (A) Ligand-receptor pair analysis of mesenchymal subpopulations with monocytes, macrophages, and osteoclasts. (B) Violin plots of osteoclast regulatory factors in mesenchymal subpopulations. (C) Violin plots of receptors for osteoclast regulatory factors in monocyte-macrophage lineage cells. (D) Representative fluorescent image of TRAP staining in 3-month-old *Adipoq/Td/Col1-GFP* mouse femur reveals that Td^+^ MALPs extend cell process contacting osteoclast. Panels b and c are the enlarged images of boxed area in a. Yellow arrows point to Td^+^ cell processes touching nearby bone surface osteoclasts (white). Note that osteoblasts (green) and osteoclasts are often located at the opposite sides of bone (c). GP: growth plate; CB: cortical bone; BM: bone marrow. Scale bar=500 μm.

Among the identified ligand-receptor pairs, the most prominent ones were RANKL-RANK and Csf1-Csf1r, two major signals for osteoclastogenesis. Violin plots clearly showed that MALPs are the major source of *Tnfsf11* and *Csf1* among mesenchymal cells (Fig. 3B). Other factors known for regulating osteoclast proliferation, migration, and differentiation, such as *Il7* (25), *Il34* (26), *Ccl2* (27), *Vcam1* (28), and *C3* (29), were also highly expressed in MALPs but not in osteoblasts and osteocytes. Their receptor expression was confirmed in monocytic cells (Fig. 3C).

Cell-cell interaction is the major mechanism by which RANKL stimulates osteoclast maturation (30, 31). In the bone marrow of *Adipoq/Td/Col1-GFP* mice, we observed that TRAP^+^, bone attaching osteoclasts are always contacted by cell processes of neighboring Td^+^ MALPs (Fig. 3Da-c). In contrast, the direct contact between osteoclasts and GFP^+^ osteoblasts was less frequent. Instead, we often observed that a line of GFP^+^ osteoblasts and a line of TRAP^+^ osteoclasts are located at the opposite sides of a trabecula (Fig. 3Dc). These data indicate that MALPs are more likely to spatially regulate osteoclastogenesis than osteoblasts via RANKL surface expression.

### RANKL from MALPs is critical for bone resorption

To study the role of RANKL in adipogenic lineage cells, we first analyzed its expression in the bone marrow of *Adipoq/Td* mice. qRT-PCR revealed that Td^+^ cells, which is only 0.74% of total bone marrow cells as analyzed by flow cytometry, express *Tnfsf11* at 15 times more than Td^−^ cells (Fig. 4A), indicating that MALPs are one of RANKL sources in the bone marrow. Next, we constructed *Adipoq-Cre Tnfsf11*^*flox/flox*^ *(RANKL CKO*^*Adipoq*^) mice. Compared to *WT* siblings, these mice had 60% and 75% decreases of *Tnfsf11* mRNA in bone marrow at 1 and 3 months of age, respectively (Fig. 4B). *Tnfsf11* mRNA in cortical bone, however, was unchanged (Fig. 4C), suggesting that these mice have RANKL deficiency specifically in adipogenic lineage cells within the marrow but not in osteocytes within the cortical bone.

**Figure 4.**
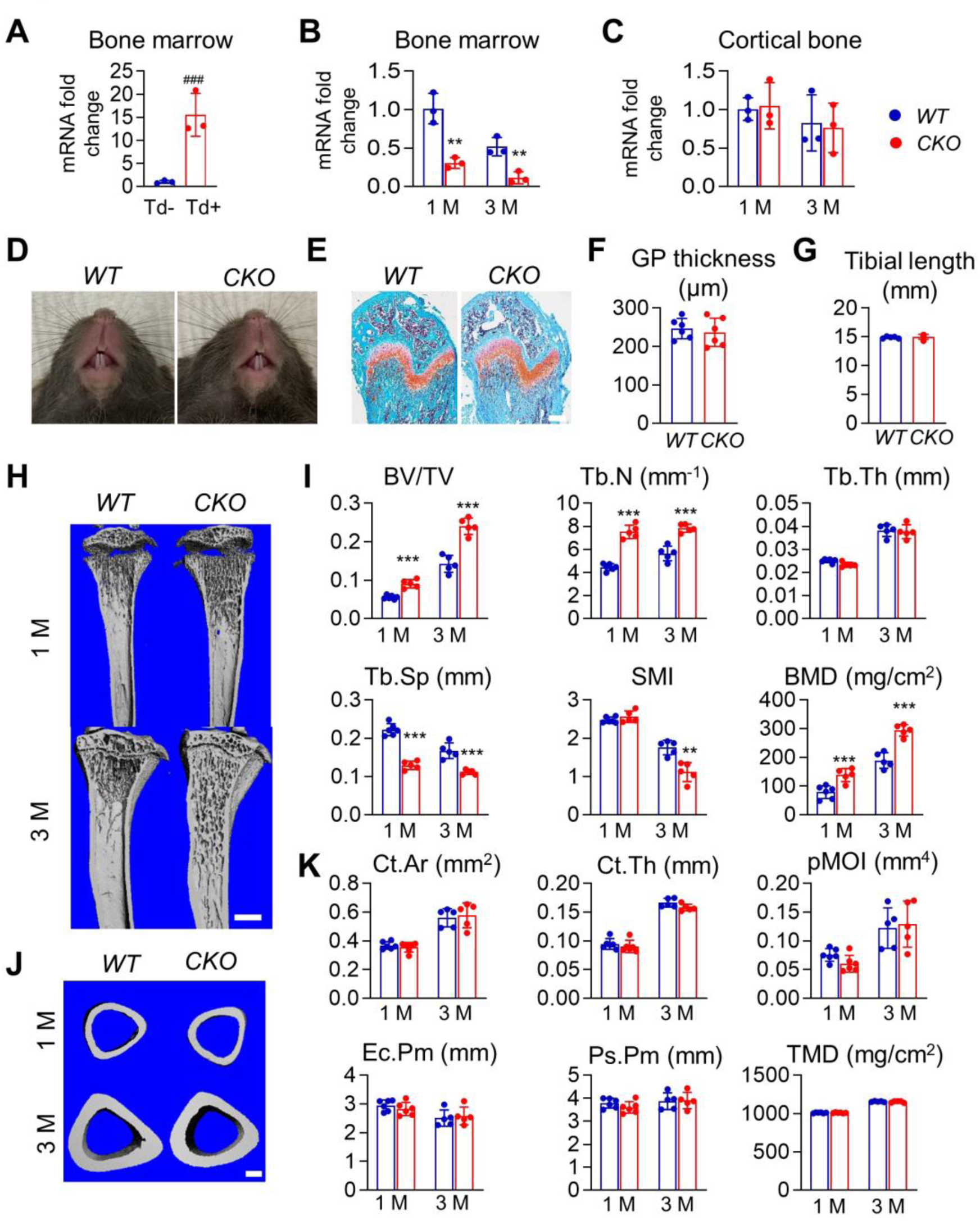
*RANKL CKO*^*Adipoq*^ mice have high trabecular bone mass. (A) qRT-PCR analysis of *Tnfsf11* mRNA in Td^+^ and Td^−^ cells sorted from bone marrow of *Adipoq/Td* mice. n=3 mice/group. (B) qRT-PCR analysis of *Tnfsf11* mRNA in bone marrow of *WT* and *RANKL CKO*^*Adipoq*^ mice at 1 and 3 months of age. n=3 mice/group. (C) qRT-PCR analysis of *Tnfsf11* mRNA in cortical bone of *WT* and *RANKL CKO*^*Adipoq*^ mice at 1 and 3 months of age. n=3 mice/group. (D) Tooth eruption is not affected in *RANKL CKO*^*Adipoq*^ mice. (E) Representative Safranin O/fast green staining of long bone sections from 1-month-old *WT* and *RANKL CKO*^*Adipoq*^ mice. Scale bar=200 μm. (F) Quantification of tibial growth plate thickness. n=6 mice/group. (G) Quantification of tibial length. n=6 mice/group. (H) 3D microCT reconstruction of *WT* and *RANKL CKO*^*Adipoq*^ mouse tibiae reveals a drastic increase of trabecular bone at 1 and 3 months of age. Scale bar=2 mm. (I) MicroCT measurement of trabecular bone structural parameters from the secondary spongiosa region. BV/TV: bone volume fraction; Tb.N: trabecular number; Tb.Th: trabecular thickness; Tb.Sp: trabecular separation; SMI: structural model index; BMD: bone mineral density. n=5-6 mice/group. (J) 3D microCT reconstructions of the midshaft region. Scale bar=0.2 mm. (K) MicroCT measurement of cortical bone structural parameters from the midshaft region. Ct.Ar: cortical area; Ct.Th: cortical thickness; pMOI: Polar moment of inertia; Ec.Pm: endosteal perimeter; Ps.Pm: periosteal perimeter; TMD: tissue mineral density. n=5-6 mice/group. ###: p<0.001 Td^+^ vs Td^−^ cells; **: p<0.01; ***: p<0.001 *CKO* vs *WT*.

*RANKL CKO*^*Adipoq*^ mice displayed normal postnatal growth with unchanged body and spleen weight up to 12 weeks of age (Fig. S3). Their tooth eruption (Fig. 4D) and growth plates (Fig. 4E, F) appeared normal with unaffected long bone growth (Fig. 4G). At 1 month of age, male *RANKL CKO*^*Adipoq*^ mice showed a marked 61% increase in tibial trabecular bone mass (BV/TV), accompanied by a 63% increase in trabecular number (Tb.N.) and a 40% decrease in trabecular separation (Tb.Sp.) (Fig. 4H, I). Trabecular thickness and structure model index (SMI) remained the same. In contrast, cortical bone was not affected with all structural parameters, such as cortical area (Ct.Ar), cortical thickness (Ct.Th), endosteal perimeter (Ec.Pm), periosteal perimeter (Ps.Pm), tissue mineral density (TMD), and MOI being the same between *CKO* and *WT* mice. At 3 months of age, the high bone mass phenotype became more striking with a 1.7-fold increase in tibial trabecular BV/TV. Cortical bone was still unaltered. Similar bone phenotypes were also observed in female mice (Fig. S4). Notably, trabecular BV/TV in adult female *CKO* mice was 2.9- and 2.0-fold higher than *WT* at tibial and vertebral sites, respectively. For comparison, we constructed *Dmp1-Cre Tnfsf11*^*flox/flox*^ *(RANKL CKO*^*Dmp1*^) mice to knockdown *Tnfsf11* expression in osteocytes. In line with previous reports (11, 13), at 1 month of age, these mice displayed a merely 18% increase in trabecular bone mass (Fig. S5). Taken together, our data indicate that MALPs contribute more to trabecular bone remodeling than osteocytes in young mice.

We next performed histology to analyze cellular changes in *RANKL CKO*^*Adipoq*^ mice. Overall, *WT* mice at 1 month of age had many more osteoclasts and osteoblasts than at 3 months of age, indicating a higher bone turnover (Fig. 5). TRAP staining revealed that osteoclast surface and number in trabecular bone of 1-month-old *RANKL CKO*^*Adipoq*^ mice are decreased by 75% and 65%, respectively, compared to *WT* mice (Fig. 5A, B). However, osteoclast formation at COJ and the endosteal surface of cortical bone was not changed, indicating that MALP-derived RANKL is not the decisive factor for osteoclastogenesis at those skeletal sites. To study whether RANKL deficiency affects osteoclast progenitors, we harvested bone marrow cells (BMCs) for in vitro osteoclastogenesis. BMCs from *WT* and *CKO* mice gave rise to the same quantity of multinucleated osteoclasts after M-CSF and RANKL induction (Fig. S6), indicating that MALP-derived RANKL does not affect the number of osteoclast progenitors.

**Figure 5.**
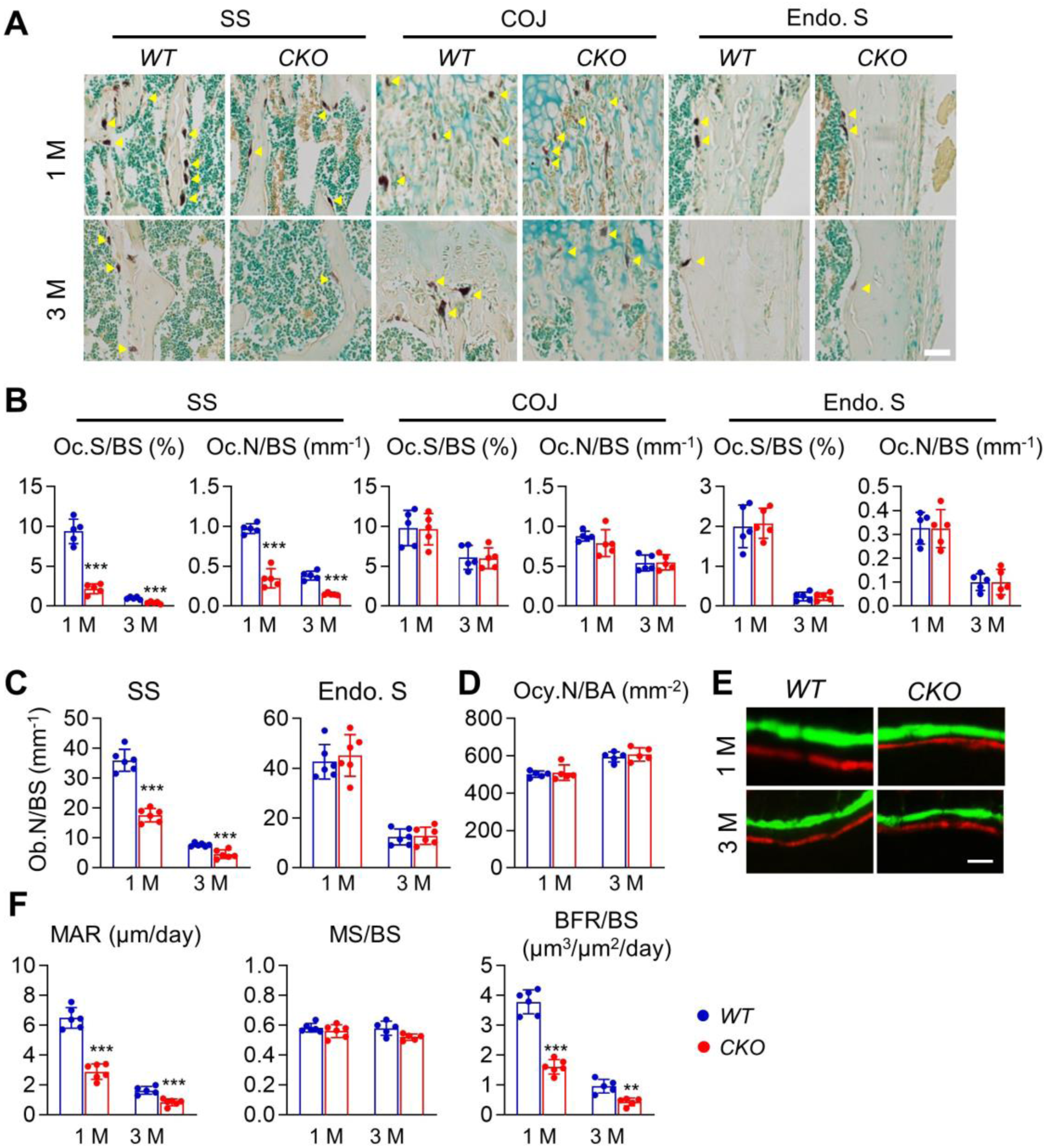
Bone resorption as well as bone formation are reduced in *RANKL CKO*^*Adipoq*^ mice. (A) Representative TRAP staining images show TRAP^+^ osteoclasts at different skeletal sites: secondary spongiosa (ss), COJ, and endosteal surface (Endo. S). Scale bar=50 μm. (B) Quantification of osteoclast surface (Oc.S/BS) and osteoclast number (Oc.N/BS) at 3 skeletal sites. BS: bone surface. n=5-6 mice/group. (C) Quantification of osteoblast number (Ob.N) in the secondary spongiosa and at the endosteal surface. n=5-6 mice/group. (D) Quantification of osteocyte density (osteocyte number per bone area, Ocy.N/BA) in the secondary spongiosa. n=5-6 mice/group. (E) Representative double labeling in distal femurs of *WT* and *CKO* mice. Scale bar=10 μm. (F) Bone formation activity is quantified. MAR: mineral apposition rate; MS: mineralizing surface; BFR: bone formation rate. n=5-6/group. **: p<0.01; ***: p<0.001 *CKO* vs *WT*.

### Suppressed bone resorption in RANKL CKO^Adipoq^ mice leads to attenuated bone formation

Meanwhile, we observed that compared to that in *WT* mice, osteoblast number is significantly reduced by 52% and 41% in the trabecular bone of 1- and 3-month-old *RANKL CKO*^*Adipoq*^ mice, respectively (Fig. 5C), while osteocyte density is not affected (Fig. 5D). Double labeling showed a decrease in osteoblast activity (Fig. 5E). Specifically, MAR and BFR were reduced by 55% and 58%, respectively, in 1-month-old *CKO* mice (Fig. 5F). Similar to osteoclasts, osteoblasts on endosteal cortical bone surface remained unchanged (Fig. 5C). These results clearly demonstrate that osteoclasts play a critical role in promoting osteoblast formation, which is consist with previous discoveries that osteoclasts “talk back” to osteoblasts, such as the reverse signaling of RANKL/RANK (32) and the forward signaling of Ephrin2/EphB4 (33).

Osteoblasts are derived from bone marrow mesenchymal progenitors. To test whether those cells are affected in *CKO* mice, we performed CFU-F assay. Strikingly, CFU-F frequency from bone marrow of 1-month-old *CKO* mice was drastically decreased by 67% (Fig. S7A). However, once seeded, their growth curve was similar to those cells from *WT* mice (Fig. S7B). Furthermore, when subjected to differentiation, these progenitors exhibited similar levels of osteogenic and adipogenic differentiation, evidenced by lineage specific staining (Fig. S7C) and marker gene expression (Fig. S7D, E). These data implicate that suppressed bone resorption in *CKO* mice reduces the pool of bone marrow mesenchymal progenitors but did not affect their proliferative and differentiation ability.

### MALP-derived RANKL contributes to pathologic bone loss

To understand the functional role of MALP-derived RANKL in osteoclast-medicated bone resorption, we tested two mouse models of pathologic bone loss. In the calvaria of *Adipoq/Td* mice, Td^+^ cells were detected abundantly inside the bone marrow but not in the suture and periosteum (Fig. 6A). All Td^+^ cells had no lipid accumulation (data not shown), indicating that they are MALPs. In the first model, we injected lipopolysaccharide (LPS) above calvaria of 6-week-old mice to induce bone loss that mimics bacteria-induced bone loss. One week later, we found a drastic increase of bone destruction in *WT* calvaria but not in *RANKL CKO*^*Adipoq*^ calvaria (Fig. 6B, C). TRAP staining revealed that LPS injections increased TRAP stained area by 18-fold and osteoclast number in *WT* calvaria by 45- and 34-fold, respectively (Fig. 6D-G). In *CKO* mice, such increases were almost completely abolished.

**Figure 6.**
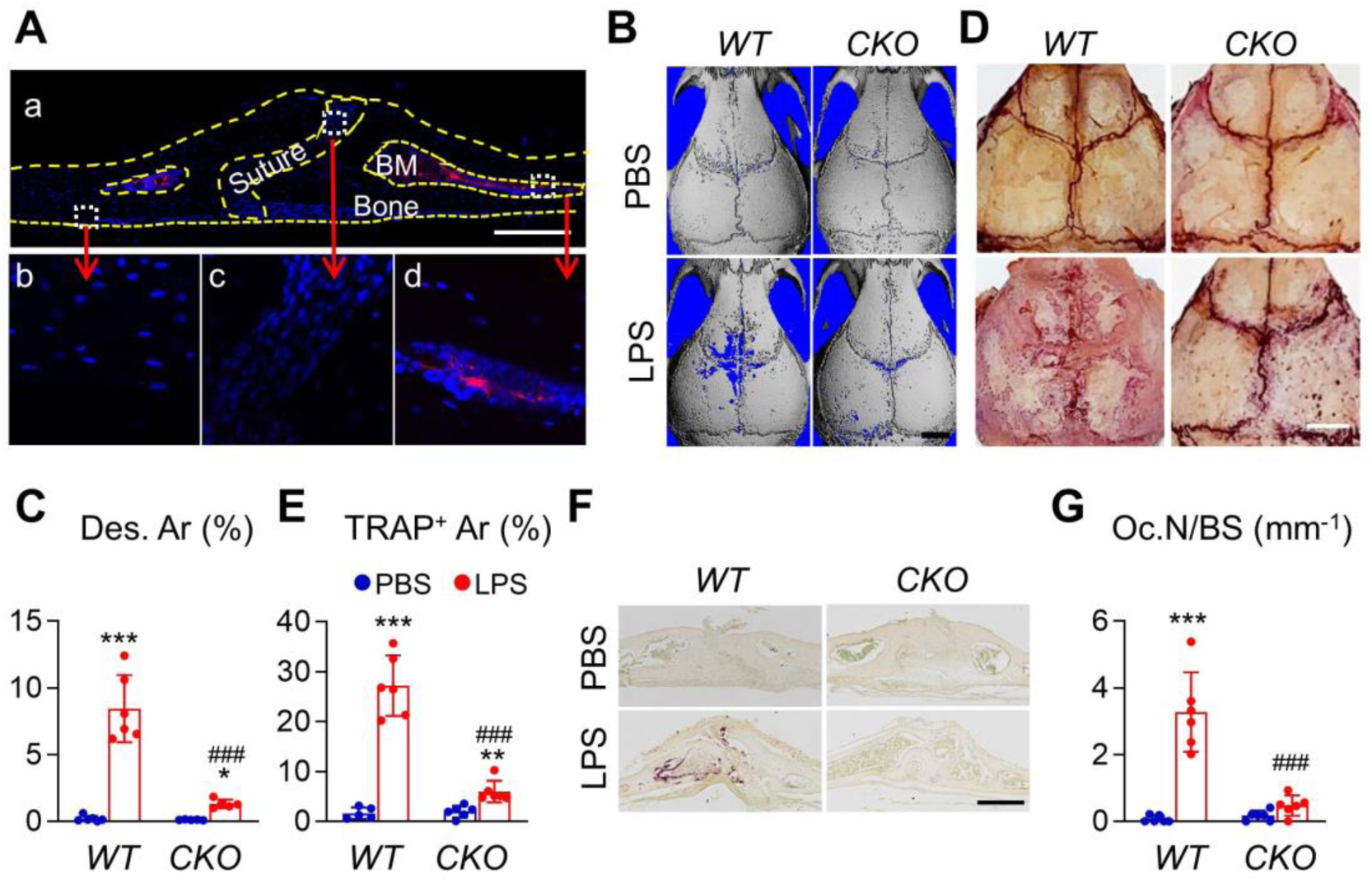
*RANKL CKO*^*Adipoq*^ mice are protected from LPS-induced calvarial bone lesions. (A) Representative coronal section of 1.5-month-old *Adipoq/Td* mouse calvaria. Bone surfaces are outlined by dashed lines. Boxed areas in the low magnification image (a) are enlarged to show periosteum (b), suture (c), and bone marrow (BM, d) regions. Scale bar=200 μm. (B) Representative 3D microCT reconstruction of mouse calvaria after 1 week of vehicle (PBS) or LPS injections. Scale bar=2 mm. (C) Quantification of percentages of bone destruction area (Des. Ar) in calvaria. n=6 mice/group. (D) Representative images of TRAP staining of whole calvaria. Scale bar=2 mm. (E) Quantification of percentages of TRAP^+^ area in calvaria. n=6 mice/group. (F) Representative images of calvaria coronal section stained by TRAP. Scale bar=2 mm. (G) Quantification of osteoclast surface (Oc.S) and number (Oc.N) in calvaria. n=6 mice/group. *: p<0.05, **: p<0.01, ***: p<0.001, LPS vs PBS; ###: p<0.001, *CKO* vs *WT*.

In the second model, we performed ovariectomy (ovx) on female *CKO* mice to mimic postmenopausal osteoporosis and examined vertebral trabecular bone 1.5 month later. Estrogen deficiency was confirmed by an 86% decrease in uterine weight in both *WT* and *CKO* mice (Fig. S8). No body weight change was observed. Ovx reduced trabecular BV/TV by 50% due to a 34% decrease in Tb.N and a 10% decrease in Tb.Th in *WT* mice (Fig. 7A, B). *CKO* mice also showed a 30% reduction in BV/TV. Interestingly, while Tb.Th was similarly decreased, Tb.N remained the same. Histology revealed that increases of osteoclast number and surface are much greater in *WT* mice (118% and 82%, respectively) than in *CKO* mice (81% and 45%, respectively, Fig. 7C, D). Meanwhile, osteoblast activity, as measured by MAR and BFR, was increased in both genotypes after ovx (Fig. 7E, F). Taken together, the above data demonstrate that RANKL from MALPs are primarily responsible for osteolytic lesions in the LPS treatment model and partially responsible for enhanced bone resorption in the ovx model.

**Figure 7.**
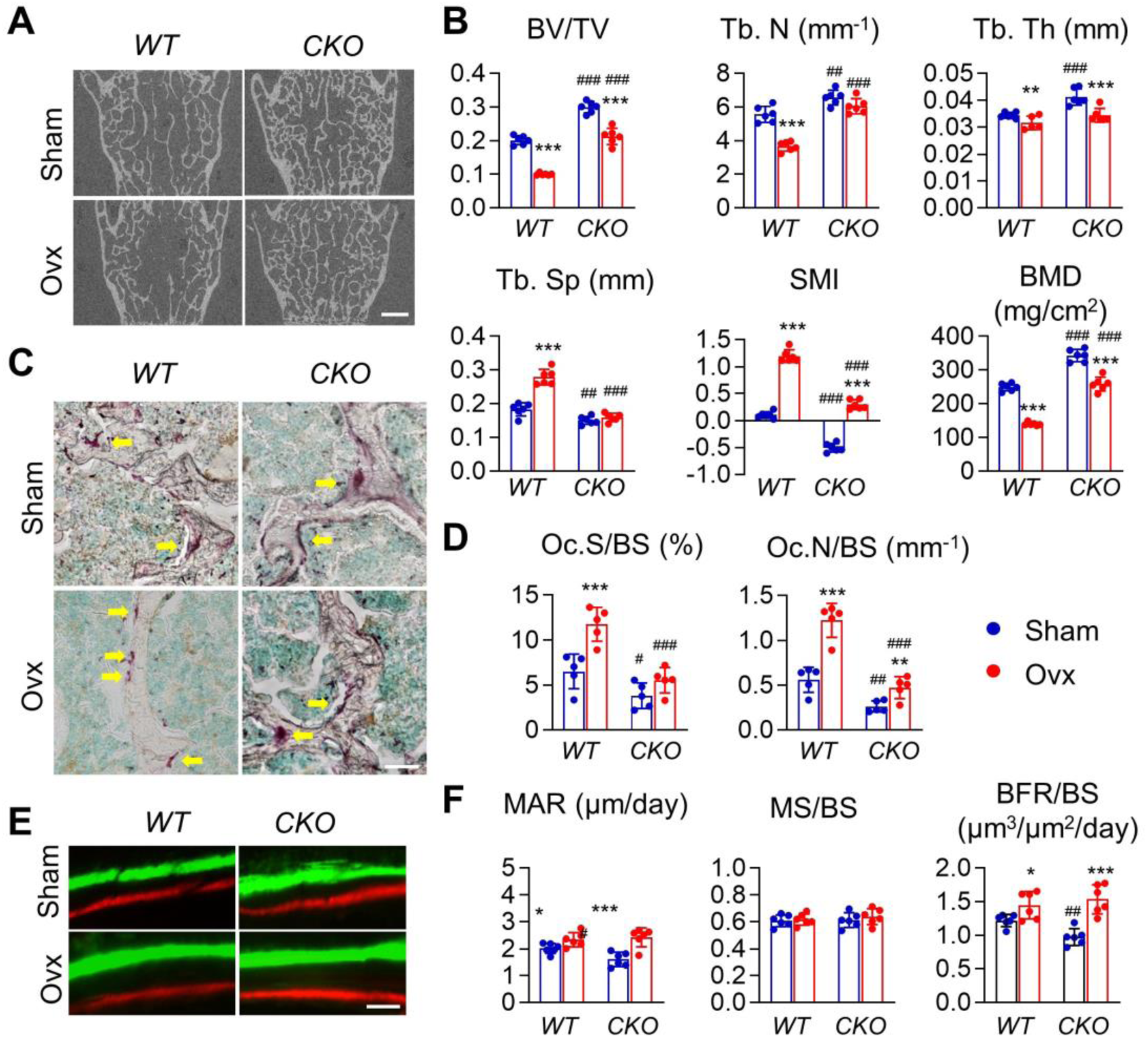
Ovx-induced bone resorption is partially attenuated in *RANKL CKO*^*Adipoq*^ mice. (A) 2D microCT reconstruction of *WT* and *RANKL CKO*^*Adipoq*^ mouse vertebrate at 1.5 months after sham or ovx surgery. Scale bar=500 μm. (B) MicroCT measurement of trabecular bone structural parameters. BV/TV: bone volume fraction; Tb.N: trabecular number; Tb.Th: trabecular thickness; Tb.Sp: trabecular separation; SMI: structural model index; BMD: bone mineral density. n=5-6 mice/group. (C) Representative image of TRAP staining in vertebrate after sham or ovx surgery. Scale bar=100 μm. (D) Quantification of osteoclast surface (Oc.S/BS) and number (Oc.N) in vertebrate after surgery. n=5-6 mice/group. (E) Representative double labeling in vertebrate of *WT* and *CKO* mice after surgery. Scale bar=10 μm. (F) Bone formation activity is quantified. MAR: mineral apposition rate; MS: mineralizing surface; BFR: bone formation rate. n=5-6 mice/group. *: p<0.05, **: p<0.01, ***: p<0.001, Ovx vs Sham; #: p<0.05, ##: p<0.01, ###: p<0.001, *CKO* vs *WT*

## Discussion

Bone remodeling, a balance between osteoblastic bone formation and osteoclastic bone resorption, is critical for the maintenance of skeletal structural integrity and mineral homeostasis. It has been long conceived that osteogenic cells are the major support cells for osteoclastogenesis, and thus they promote bone resorption. In this work, we demonstrated that the committed adipogenic precursors, MALPs, are another important player in controlling bone resorption. In silico analysis revealed that they have the most interactions with monocyte-macrophage lineage cells among mesenchymal subpopulations and predicted that they express critical osteoclast regulator factors, including RANKL, at a much higher level than other mesenchymal cells, including osteoblasts and osteocytes, in young and adult mice. Strikingly, adipogenic specific knockdown of RANKL causes a drastic high bone mass phenotype (60-70% increase in BV/TV) as early as 1 month of age. In contrast, at the same age, mice with osteocyte-specific deficiency of RANKL showed only small bone changes in our hands (an 18% increase in BV/TV) or no changes in other groups (11, 13). Furthermore, MALP-derived RANKL is absolutely required for LPS-induced osteolysis and partially required for ovx-induced bone loss, reinforcing the importance of MALPs as an important cellular regulator of bone remodeling under normal and diseased conditions.

Past studies have provided mostly in vitro evidence that bone marrow adipocytes support osteoclast formation. Using an elegant inverted coculture method, Goto et al. showed that primary human bone marrow LiLAs stimulate TRAP^+^ multinucleated osteoclast formation in the presence of TNFα or dexamethasone via upregulation of RANKL (34, 35). Adipogenic differentiation of mouse bone marrow mesenchymal progenitors was found to be associated with increased expression of RANKL and decreased expression of OPG, a decoy receptor of RANKL (36). Further studies showed that adipogenic TFs, C/EBPβ and δ, activate RANKL gene transcription. Interestingly, mesenchymal progenitors from aged mice are better at supporting osteoclast formation in coculture than those from young mice when utilizing adipogenic differentiation medium. This is consistent with the well-known effects of aging on bone marrow adiposity and our finding that aging expands the MALP population (14). A recent study carefully dissected adipocytes derived from bone marrow mesenchymal progenitors in culture into non-lipid-laden and lipid-laden ones, which bear resemblance to MALPs and LiLAs in our analysis, respectively. A striking finding was that RANKL is mostly presented in non-lipid-laden adipocytes but not in lipid-laden adipocytes (37). Since scRNA-seq cannot capture LiLAs, we do not know whether *Tnfsf11* expression is down-regulated when MALPs become LiLAs. However, these in vitro data as well as the high ratio of MALPs vs LiLAs in vivo strongly indicate that MALPs are the major cell type controlling bone resorption in vivo. Further, we provide the first in vivo evidence that MALP-derived RANKL controls trabecular bone remodeling.

Marrow adipose tissue (MAT) is a unique adipose tissue that is morphologically and functionally distinct from peripheral adipose tissues (38). Traditionally, it has only referred to LiLAs as in other adipose tissues. Our discovery of MALPs adds another important and abundant cell population to this tissue although no counterpart of MALPs has been identified in other adipose tissues. Hence, it is not surprising that MAT possesses a set of functions that does not seem to exist in other adipose tissues. Our previous study revealed that MALPs maintain bone marrow vessel integrity and inhibit bone formation (14). Here we show that MALPs promote bone resorption. In line with our results, it was reported that only bone marrow adipocytes but not peripheral adipocytes express RANKL (39). The unique functionality attributable to MAT is likely due to its special location in bone marrow, where bone remodeling and hematopoiesis occur constantly. Having a cell body and multiple processes, MALPs form a 3D network structure that contacts almost every cell inside the bone. Hence, we propose that their main function is to regulate the bone marrow environment, including bone resorption.

While our single cell datasets and subsequently animal studies identify MALPs as the major support cells for osteoclastogenesis, we cannot rule out the importance of other cells. One limitation of our single cell approach is that our dataset might contain only young osteocytes but not mature ones because Td^+^ cells were collected via enzymatic digestion of bones longitudinally cut into half after flushing out bone marrow. Since *Tnfsf11* mRNA is more than 10 times higher in osteocytes than in osteoblasts (12), it is likely that mature osteocytes play a more important role in controlling osteoclastogenesis than young, surface osteocytes. Interestingly, *RANKL CKO*^*Adipoq*^ mice displayed reduced bone resorption only within trabecular bone but not at cortical bone. Since MALPs do not exist at periosteum, it is expected that periosteal bone resorption is not affected in CKO mice. However, MALPs are abundant at the endosteal surface. To reconcile these data, we propose that mature osteocytes, which are abundant in cortical bone but relatively scarce in trabecular bone, control osteoclast formation at the surface of cortical bone. In line with this idea, Xiong et al. found that *RANKL CKO*^*Dmp1*^ mice are resistant to tail suspension-induced cortical bone loss (11). Similarly, MALP-derived RANKL does not contribute to cartilage-to-bone remodeling during endochondral ossification even MALPs are abundant at COJ. Hypertrophic chondrocytes control osteoclast formation in this event (11). Moreover, during estrogen deficiency, RANKL produced from B cells and osteocytes, in addition to MALPs, is required for the enhanced osteoclastic activity (40, 41). Our studies also demonstrate a dominant action of MALPs in LPS-induced osteolysis in calvaria. Collectively, osteoclast formation is controlled by a variety of cells in bone in skeletal site-specific and disease-dependent manners (Figure 8). Since *Dmp1-Cre* is somewhat leaky and expressed in osteoblasts (42), we believe that in the trabecular bone of adult mice, MALPs, osteocytes, and osteoblasts all contribute to promoting bone resorption by osteoclasts.

**Figure 8.**
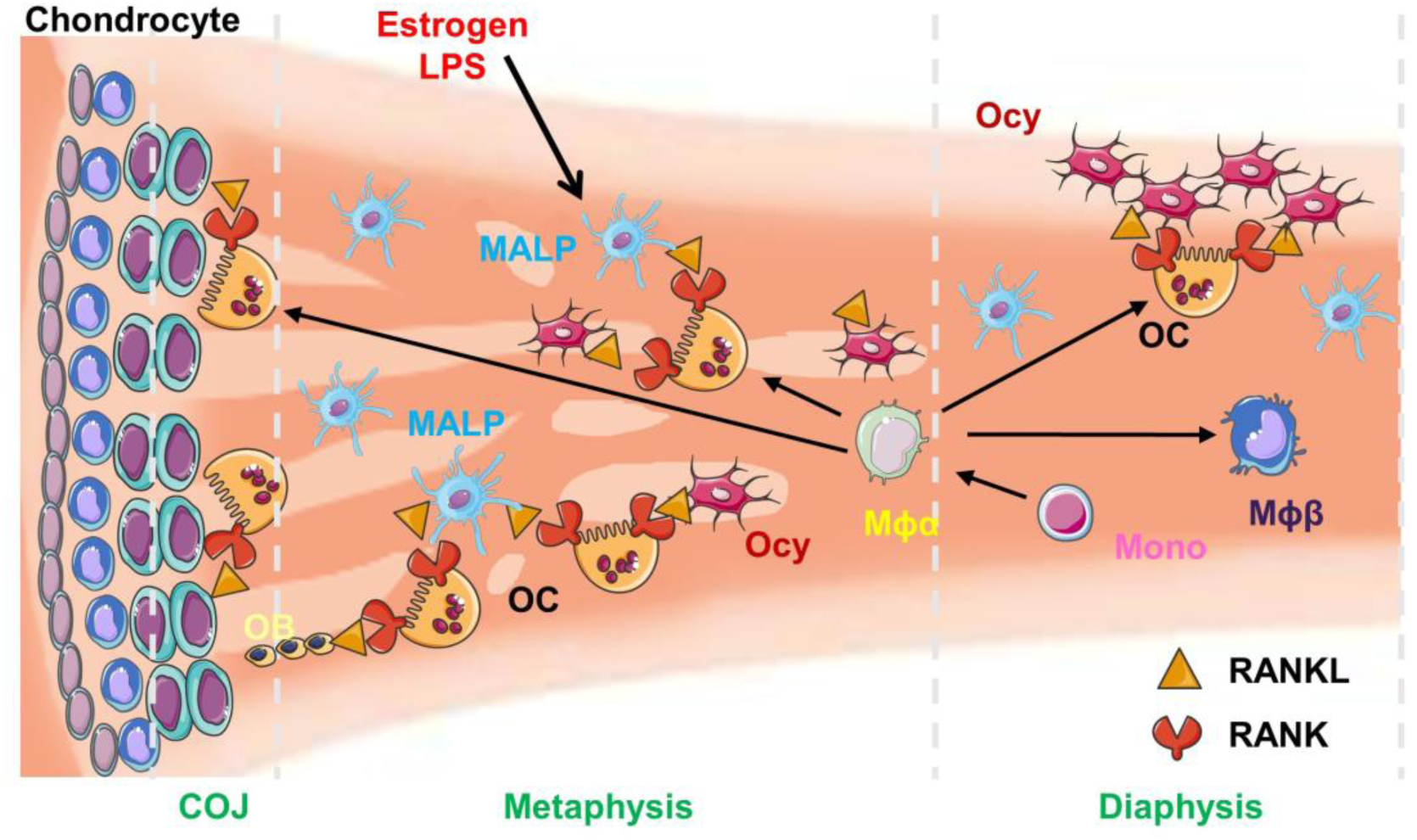
A schematic diagram depicts in vivo osteoclast differentiation and its regulation in a site-specific manner. In bone marrow, Mϕα, derived from either embryonic macrophages or bone marrow monocytes, undergoes bi-lineage differentiation into osteoclasts and Mϕβ. At COJ, RANKL from hypertrophic chondrocytes stimulates osteoclast formation. At the endosteal surface of cortical bone, osteocytes are the major producer of RANKL for osteoclastogenesis. Within the trabecular bone, MALPs, osteocytes, and possibly osteoblasts, all promote osteoclast differentiation. In young mice, RANKL from MALPs seems to play a predominant role in this process. External stimuli, such as estrogen and LPS, could act on MALPs to stimulate bone resorption. OC: osteoclast; OB: osteoblast; Ocy: osteocyte; Mono: monocyte; Mϕ: macrophage.

It would be interesting to explore the action of MALPs in other bone disorders and therapies. Parathyroid hormone (PTH1-34) is an FDA-approved drug for improving bone mass in osteoporosis patients (43). Ablation of its receptor in skeletal mesenchymal progenitors using *Prx1-Cre* leads to increased RANKL expression in bone marrow adipocytes, enhanced marrow adiposity, and bone resorption (39), implying a role of MALPs in the anabolic actions of PTH.

Our scRNA-seq dataset and analysis have another limitation. Since our original purpose was to isolate mesenchymal lineage cells, the *Col2/Td* mouse model was adopted. However, for reasons we do not yet understand, cell sorting of Td^+^ cells from these mice includes a small portion of hematopoietic cells and endothelial cells, a fact that we used to our advantage for the subsequent monocyte-macrophage lineage and cell-cell interaction analyses. However, we cannot exclude the possibility that hematopoietic cells in our dataset do not contain all subsets of monocytes, macrophages, and osteoclasts. Therefore, the pseudotime trajectory analysis might not be inclusive. Moreover, recent studies have demonstrated that tissue-resident macrophages contain both embryonic stage-derived macrophages and adult HSC-derived macrophages (44) and that osteoclasts in adult bone are also fusion of cells originated from these two types of macrophages (45, 46), raising a concern whether scRNA-seq analysis could distinguish these two sources of progenitors. Nevertheless, our analysis revealed that Mϕα cells differentiate into two types of terminal cells, osteoclasts that eat bone matrix and Mϕβ that eat apoptotic cells in bone. Mϕα cells could be a combination of embryonic stage-derived and HSC-derived macrophages. Interestingly, they express a variety of cytokines and chemokines, in accordance with previous findings that bone marrow macrophages regulate hematopoiesis and bone formation (47).

The current mainstay drugs for osteoporosis are anti-resorptive agents, such as bisphosphonates and anti-RANKL antibody (denosumab) (48). However, their long term use has caused concern for undesired effects, such as atypical fracture and osteonecrosis. Therefore, identifying a cell population that can regulate both osteoblasts and osteoclasts represents a clinically prudent approaches that will allow for the fine-tuning of bone remodeling not only for better overall efficacy but also eventually for precise individualized effect. MALPs not only produce RANKL but also other adipokines and cytokines that have osteoclast regulatory actions. Furthermore, ablation of this cell population causes the most profound bone formation we have ever observed, indicating a pivotal role in regulating bone generation and regeneration (14). Further strategies to understand the mechanisms of the dual actions of MALPs on bone remodeling and seeking approaches to therapeutically target this cell population could be of critical value in developing new treatment for osteoporosis and other disorders of pathologic bone loss.

## Author contributions

W.Y. and L.Q. designed the study. W.Y. and Y.W. performed animal experiments. L.Y. helped with library construction. W.Y., L.Z. and T.G. performed histology and imaging analysis. W.Y., Z.L., and L.Z. performed cell culture and qRT-PCR experiments. W.Y., L.Y. and L.Q. performed computational analyses. H.K., N.D., X.S., S.Y., Y.C. and J.A. provided administrative, technical, or material support and consultation. L.Q. wrote the manuscript. W.Y., L.Z., L.Y., N.D., S.Y. Y.C. and J.A. reviewed and revised the manuscript. L.Q. approved the final version.

## Acknowledgments

This study was supported by NIH grants NIH/NIAMS R01AR066098, R21AR074570 (to L.Q.), R00AR067283 (to N.D.), R01AR 066101 (to S.Y.), AR0569546 (to Y.C.) and P30AR069619 (to Penn Center for Musculoskeletal Disorders)

