## supplementary for "Bone marrow adipogenic lineage precursors (MALPs) promote osteoclastogenesis in bone remodeling and pathologic bone loss"

### Supplementary Methods

*Animals study design*

All animal work performed in this report was approved by the Institutional Animal Care and Use Committee (IACUC) at the University of Pennsylvania. In accordance with the standards for animal housing, mice were group housed at 23-25˚C with a 12 h light/dark cycle and allowed free access to water and standard laboratory pellets.

*Col2-Cre Rosa-tdTomato* (*Col2/Td*), *Adipoq-Cre Rosa-tdTomato* (*Adipoq/Td*) mice were generated by breeding *Rosa-tdTomato* (Jackson Laboratory, Bar Harbor, ME, USA) mice with *Col2-Cre* (49) and *Adipoq-Cre* (50) mice respectively. *Adipoq-Cre Rosa-tdTomato 2.3kbCol1-GFP (Adipoq/Td/Col1-GFP)* were generated by breeding *Adipoq/Td mice* with *2.3kbCol1-GFP* mice (24). *Tnfsf11*$^{flox/flox}$ mice were constructed by flanking the exon 5 of *Tnfsf11* gene with loxP sites (H.K., Y.C., unpublished data). To generate *RANKL CKO*$^{Adipoq}$ mice, we first bred *Adipoq-Cre* with *Tnfsf11*$^{flox/flox}$ mice to obtain *Adipoq-Cre Tnfsf11*$^{flox/+}$, which were then crossed with *Tnfsf11*$^{flox/flox}$ to generate *RANKL CKO*$^{Adipoq}$ mice and *WT* (*Tnfsf11*$^{flox/flox}$ and *Tnfsf11*$^{flox/+}$) siblings. *RANKL CKO*$^{Dmp1}$ mice were generated using a similar breeding strategy with *Dmp1-Cre* (51).

For LPS-induced bone destruction, 6-week-old male mice were injected with 25 mg/kg LPS (Sigma Aldrich, St. Louis, MO) or PBS above calvaria. After 7 days, calvariae were collected and analyzed by microCT followed by TRAP staining.

For ovx-induced bone destruction, ovx or sham operation was performed on 3-month-old female mice. Six weeks later, L4 and L5 vertebrae were collected and analyzed by microCT followed by dynamic histomorphometry and TRAP staining.

*Single-cell RNA sequencing of endosteal bone marrow cells*

We constructed 3 batches of single cell libraries of endosteal $Td^+$ bone marrow cells from 1-month-old (n=2), 1.5-month-old (n=3), and 3-month-old (n=3) male *Col2/Td* mice for sequencing as described previously (14). 20,000 cells were loaded in aim of acquiring one single library of 10,000 cell for each age group by Chromium controller (V3 chemistry version, 10X Genomics Inc, San Francisco, USA), barcoded and purified as described by the manufacturer, and sequenced using a 2x150 pair-end configuration on an Illumina HiSeq platform at a sequencing depth of ~400 million reads. Cell ranger (Version 3.0.2, https://support.10xgenomics.com/single-cell-geneexpression/software/pipelines/latest/what-is-cell-ranger) was used to demultiplex reads, followed by extraction of cell barcode and unique molecular identifiers (UMIs). The cDNA insert was aligned to a modified reference mouse genome (mm10).

Seurat package V3 was used for filtering, variable gene selection, dimensionality reduction analysis and clustering standardly (52). Doublets or cells with poor quality (genes>6000, genes<200, or >5% genes mapping to mitochondrial genome) were excluded. Expression was natural log transformed and normalized for scaling the sequencing depth to a total of $1x10^4$ molecules per cell. For the integrated dataset, anchors from different dataset were defined using the FindIntegrationAnchors function, and these anchors were then used to integrate datasets together with IntegrateData. Datasets were scaled by regressing out the number of UMIs and percent mitochondrial genes. Statistically significant principle components were selected as input for uniform manifold approximation and projection (UMAP) plots. Different resolutions for clustering were used to demonstrate the robustness of clusters. In addition, differentially expressed genes within each cluster relative to the remaining clusters were identified using FindMarkers within Seurat. Sub-clustering was performed by isolating the monocytic lineage clusters using known marker genes, followed by reanalysis as described above. Cell-cycle analysis were calculated by using the Seurat Cell-cycle scoring function, proliferative cells were defined as cells in G2M or S Phase.

To computationally delineate the developmental progression of monocyte, macrophage and osteoclast and order them in pseudotime, we used the algorithms implemented in the Monocle 2 package. We include these cells for the analysis. We ordered cells by selecting genes with high dispersion across cells, using a parameter of "mean_expression >= 0.05 & dispersion_empirical >= 1 * dispersion_fit", lists of genes were selected for dimensional reduction to generate the trajectory reconstruction using the nonlinear reconstruction algorithm DDRTree. Branched expression analysis modeling, or BEAM was used to determine the genes that are differentially expressed between branches. A mouse transcript factors (TF) list (TFdb) were used to detect 64 TFs among the differential expressed genes (53).

We also performed the trajectory analysis using Slingshot. Briefly, UMAP was used as dimensional reduction after the PCA were calculated for individual or integrated datasets. Then Seurat objects were transformed into SingleCellExperiment objects. Slingshot trajectory analysis were conducted using the Seurat clustering information and with dimensionality reduction produced by UMAP.

Cluster specific markers for each cluster, relative to the remaining population, were conducted using the FindMarkers to identify differentially expressed genes (DEGs). Sub-clustering was performed by isolating the monocytic lineage clusters identified from the remaining bone marrow cells using known marker genes, followed by reanalysis as described above. DEGs between these clusters were generated as described above. GO terms and clusters, as well as

KEGG pathway enrichment, were identified using the database for annotation, visualization and integrated discovery (DAVID) (54).

To analyze cell–cell communication mediators among different cell types, we used CellPhoneDB, a repository of ligands, receptors and interaction data that relies on public information to annotate receptors and ligands (55). Briefly, each cell type cluster defined by Seurat was used as input into CellphoneDB that is based on statistical methods and analyses to generate interaction numbers of ligand/receptor pair amongst different groups.

*Micro-computed tomography (microCT) analysis*

MicroCT analysis (microCT 35, Scanco Medical AG, Brüttisellen, Switzerland) was performed at 6 μm isotropic voxel size as described previously (18). Briefly, the proximal end of the tibia corresponding to a 0 to 2.8 mm region below the growth plate was scanned at 6 μm isotropic voxel size to acquire a total of 462 μCT slices per scan. The images of the secondary spongiosa regions 0.6 to 1.8 mm below the lowest point of the growth plate were contoured for trabecular bone analysis. At the tibia midshaft, a total of 100 slices located 4.8-5.4 mm away from the proximal growth plate were acquired for cortical bone analyses by visually drawing the volume of interest (VOI). In vertebrae, the region (total about 300 slices) 50 slices away from the top and bottom end plates was acquired for trabecular bone analysis. The trabecular bone tissue within the VOI was segmented from soft tissue using a threshold of 487.0 mgHA/cm3 and a Gaussian noise filter (sigma=1.2, support=2.0). The cortical bone tissue was using a threshold of 661.6 mgHA/cm3 and a Gaussian noise filter (sigma=1.2, support=2.0). Three-dimensional standard microstructural analysis was performed to determine the geometric trabecular bone volume fraction (BV/TV), bone mineral density (BMD), trabecular thickness (Tb.Th), trabecular separation (Tb.Sp), trabecular number (Tb.N), and structure model index (SMI). For analysis of cortical bone, periosteal perimeter (Ps.Pm), endosteal perimeter (Ec.Pm), cortical bone area (Ct.Ar), cortical thickness (Ct.Th), polar moment of inertia (pMOI), and tissue mineral density (TMD) were recorded. All calculations were performed based on 3D standard microstructural analysis (56).

Calvaria microCT analysis was performed at 15 μm isotropic voxel size. The three-dimensional images were reconstructed to visualize the destructive area. A square region of 8x 8 mm centered at the midline suture was selected for further quantitative analysis by ImageJ.

*Histology and bone histomorphometry*

To obtain whole mount sections for immunofluorescent imaging of *Adipoq/Td/Col1-GFP* mouse bones, freshly dissected femurs were fixed in 4% PFA for 1 day, decalcified in 10% EDTA for 4-5 days, and then immersed into 20% sucrose and 2% polyvinylpyrrolidone (PVP) at 4°C overnight. Then sample was embedded into medium containing 8% gelatin, 20% sucrose and 2% PVP and sectioned at 50 μm in thickness. Sections were incubated with rat anti-CD45 (Biolegend, 103101), rat anti-Endomucin (Santa cruz, sc-65495), or rabbit anti-Perilipin (Cell signaling, 9349) at 4°C overnight followed by Alexa Fluor 647 anti-rat (Abcam, ab150155) or anti-rabbit (Abcam, ab150157) secondary antibodies incubation 1 hour at RT. For EdU staining, mice received 1.6 mg/kg EdU 1 day and 3 hr before sacrifice and the staining was carried out according to the manufacturer's instructions (ThermoFisher scientific, Click-iT™ EdU Alexa Fluor™ 647 Imaging Kit, D3822).

To obtain paraffin sections, mouse bones were fixed in 4% PFA for 24 hr and decalcified in a 10% EDTA for 3-4 weeks for long bones and vertebrates and 5-7 days for calvariae at 4°C. Samples were then embedded in paraffin, sectioned at 6 μm, and processed for hematoxylin and eosin (H&E) staining, Safranin O/fast green staining, or tartrate-resistant acid phosphatase (TRAP) staining using a kit (Sigma-Aldrich, 387A).

To obtain cryosections, mouse bones were dissected and fixed in 4% PFA for 24 hr, dehydrated in 30% sucrose in PBS, embedded in optimal cutting temperature (OCT) compound, and sectioned at 6 μm using a cryofilm tape (Section Lab, Hiroshima, Japan). Fluorescent TRAP staining was performed as described previously (57). Briefly, sections were incubated in TRAP buffer (0.92% sodium acetate anhydrous, 1.14% L-(+)tartaric acid, 1% glacial acetic acid, pH 4.1~4.3) for 15 minutes followed by ELF97 substrate (Life Tech, E6589) diluted at 1:40 for 5 minutes. The ELF97 substrate generates a yellow fluorescent signal when cleaved by TRAP. For dynamic histomorphometry, mice received calcein (15 mg/kg, Sigma Aldrich) at 9 (for adult mice) or 4 days (for adolescent mice) before euthanization and xylenol orange (90 mg/kg, Sigma Aldrich) at 2 days before euthanization. Sagittal cryosections of tibiae prepared with cryofilm tape were used for dynamic histomorphometry. Sections were scanned by a Nikon Eclipse 90i fiuorescence microscope and areas within secondary spongiosa were quantified by Bioquant Osteo Software (Bioquant Image Analysis, Nashville, TN, USA). The primary indices include total tissue area (TV), trabecular bone perimeter (BS), single- and double-labeled surface, and interlabel width. Mineralizing surface (MS) and surface-referent bone formation rate (BFR/BS, $\mu m^3/\mu m^2/d$) were calculated as described by Dempster et al. (58).

*Cell culture*

For CFU-F assay, flushed bone marrow cells were plated at $1x10^6$ cells/T25 flask. Cells were cultured in growth medium (α-MEM supplemented with 15% FBS, 0.1% β-mercaptoethanol, 20 mM glutamine, 100 IU/ml penicillin, and 100 µg/ml streptomycin) for 7 days before counting CFU-F number.

Mesenchymal progenitors were obtained by culturing bone marrow cells at a high density ($3x10^6$ cells/T25 flask). Once confluent, cells were switched to either adipogenic medium (DMEM with 10% FBS, 10 ng/ml triiodothyronine, 1 µM rosiglitazone, 1 µM dexamethasone, 10 µg/ml insulin, 100 IU/ml penicillin, and 100 µg/ml streptomycin) for 7 days followed by Oil Red O staining or osteogenic medium (αMEM with 10% FBS, 10 nM dexamethasone, 10 mM β-glycerophosphate, 50 µg/mL ascorbic acid, 100 IU/mL penicillin, and 100 µg/mL streptomycin) for 2 weeks followed by Alizarin Red staining. Brightfield and fluorescent images of mesenchymal progenitors from 3-month-old *Adipoq/Td* mice undergoing adipogenic and osteogenic differentiation were taken by fluorescence inverted microscopy (Nikon Eclipse, TE2000-U).

For in vitro osteoclastogenesis, bone marrow macrophages (BMMs) were obtained from the tibiae and femurs as described previously (59, 60). They were seeded at $2 \times 10^6$ cells/well in 24-well plates and stimulated with 100 ng/mL RANKL and 20 ng/mL M-CSF (R&D Systems, Minneapolis, MN, USA) for 5 days to generate mature OCs. TRAP staining was performed using a TRAP kit (Sigma Aldrich, 387A). Osteoclasts were quantified by counting the number of $TRAP^+$, multinucleated cells (≥ 3nuclei/cell) per well.

*RNA analysis*

To quantify the expression level of marker genes, total RNA was collected in Tri Reagent (Sigma Aldrich) for RNA purification. A Taqman Reverse Transcription Kit (Applied BioSystems, Inc., Foster City, CA, USA) was used to reverse transcribe mRNA into cDNA. The power SYBR Green PCR Master Mix Kit (Applied BioSystems, Inc) was used for quantitative real-time PCR (qRT-PCR). The primer sequences for the genes used in this study are listed in Supplemental Table S3.

*Statistical analyses*

Data are expressed as means ± standard deviation (SD) and analyzed by t-test, one-way ANOVA with Dunnett's or Turkey's posttest and two-way ANOVA with Turkey's post-test for multiple comparisons using Prism 8 software (GraphPad Software, San Diego, CA). For assays using primary cells, experiments were repeated independently at least three times and representative data were shown here. Values of $p<0.05$ were considered statistically significant.

### Supplementary Figure Legend

**Figure S1**

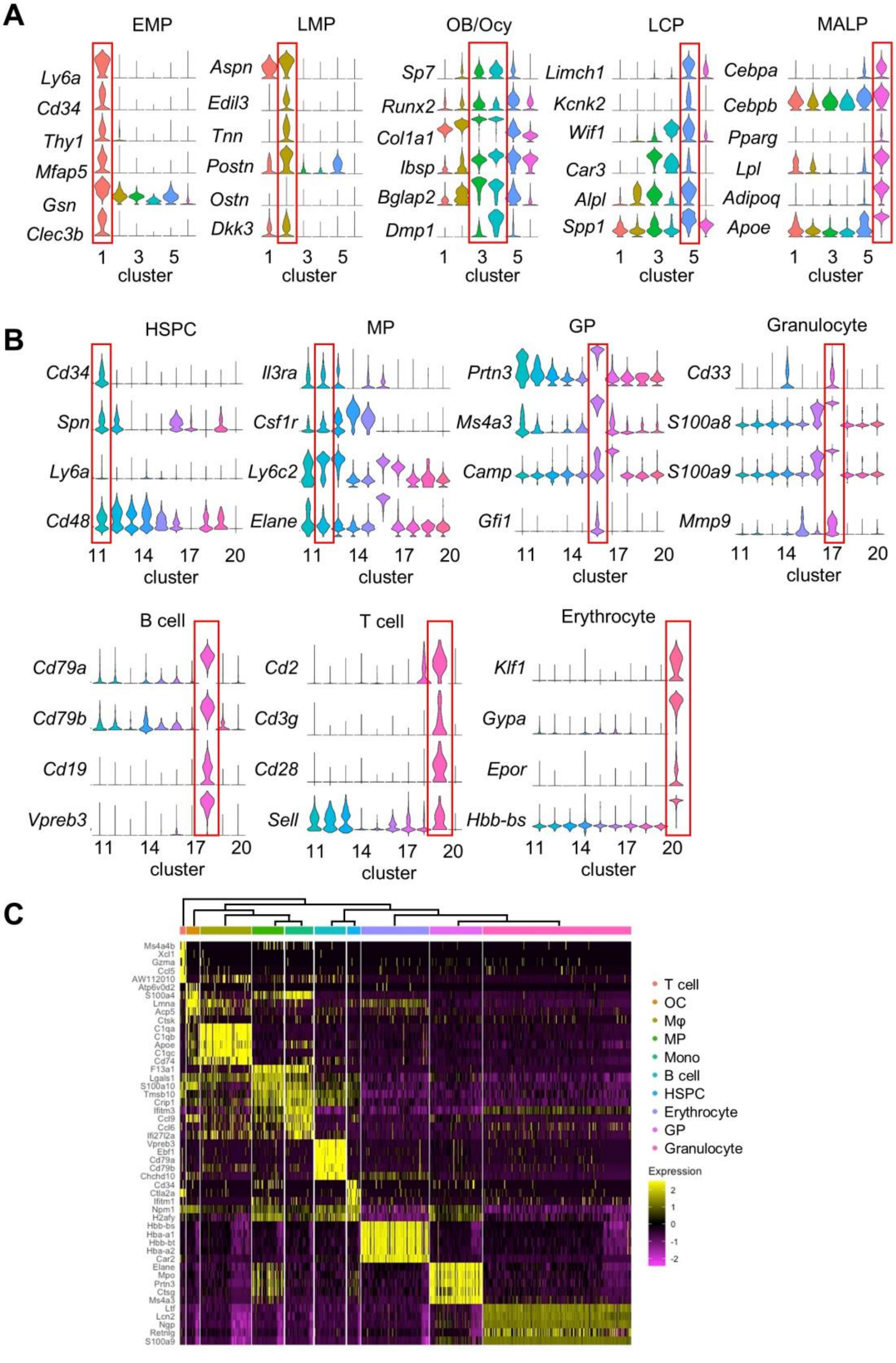
A
EMP
Ly6a
Cd34
Thy1
Mfap5
Gsn
Clec3b
LMP
Aspn
Edil3
Tnn
Postn
Ostn
Dkk3
OB/Ocy
Sp7
Runx2
Col1a1
Ibsp
Bglap2
Dmp1
LCP
Limch1
Kcnk2
Wif1
Car3
Alpl
Spp1
MALP
Cebpa
Cebpb
Pparg
Lpl
Adipoq
Apoe
1 3 5
cluster
B
HSPC
Cd34
Spn
Ly6a
Cd48
MP
Il3ra
Csf1r
Ly6c2
Elane
GP
Prtn3
Ms4a3
Camp
Gfi1
Granulocyte
Cd33
S100a8
S100a9
Mmp9
11 14 17 20
cluster
B cell
Cd79a
Cd79b
Cd19
Vpreb3
T cell
Cd2
Cd3g
Cd28
Sell
Erythrocyte
Klf1
Gypa
Epor
Hbb-bs
C
Ms4a4b
Xcl1
Gzma
Ccl5
AW112010
Atp6v0d2
S100a4
Lmna
Acp5
Ctsk
C1qa
C1qb
Apoe
C1qc
Cd74
F13a1
Lgals1
S100a10
Tmsb10
Crip1
Ifitm3
Ccl9
Ccl6
Ifi27l2a
Vpreb3
Ebf1
Cd79a
Cd79b
Chchd10
Cd34
Ctla2a
Ifitm1
Npm1
H2afy
Hbb-bs
Hba-a1
Hbb-bt
Hba-a2
Car2
Elane
Mpo
Prtn3
Ctsg
Ms4a3
Ltf
Lcn2
Ngp
Retnlg
S100a9
T cell
OC
Mφ
MP
Mono
B cell
HSPC
Erythrocyte
GP
Granulocyte
Expression
2
1
0
-1
-2

Figure S1. ScRNA-seq analysis of mesenchymal lineage and hematopoietic lineage cell clusters.

(A) Violin plots of marker gene expression in mesenchymal lineage subpopulations. EMP: early mesenchymal progenitor; LMP: late mesenchymal progenitor; OB: osteoblast; Ocy: osteocyte; LCP: lineage committed progenitor.

(B) Violin plots of marker gene expression in hematopoietic lineage subpopulations. HSPC: hematopoietic stem and progenitor cells; MP: monocyte progenitor; GP: granulocyte progenitor.

(C) Hierarchy heatmap of hematopoietic lineage cell clusters.

**Figure S2**

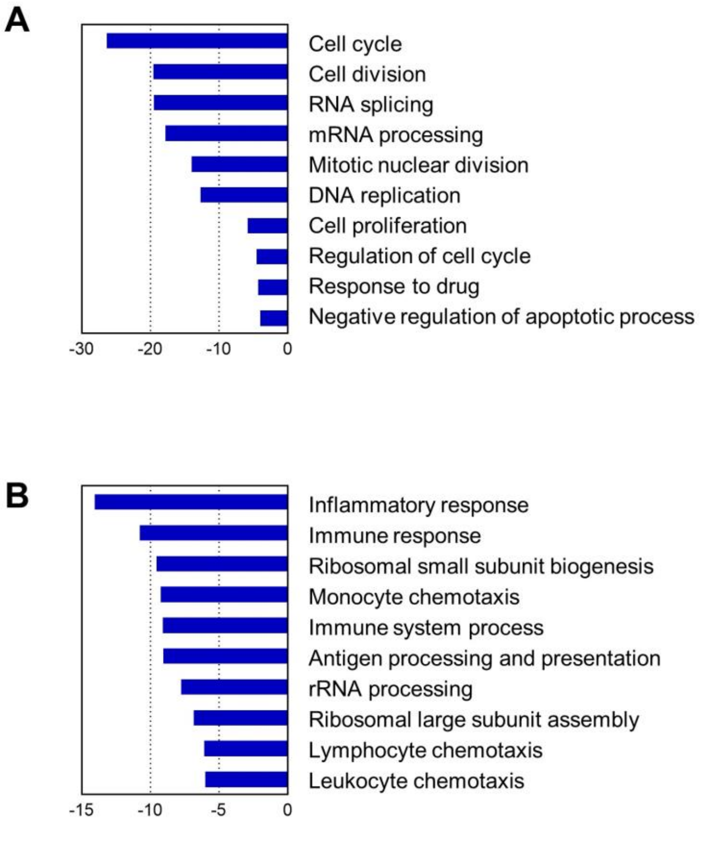


Figure S2. GO term and KEGG pathway analyses of up-regulated genes in early osteoclasts versus late osteoclasts (A) and in Mϕα versus Mϕβ cells (B).

**Figure S3**

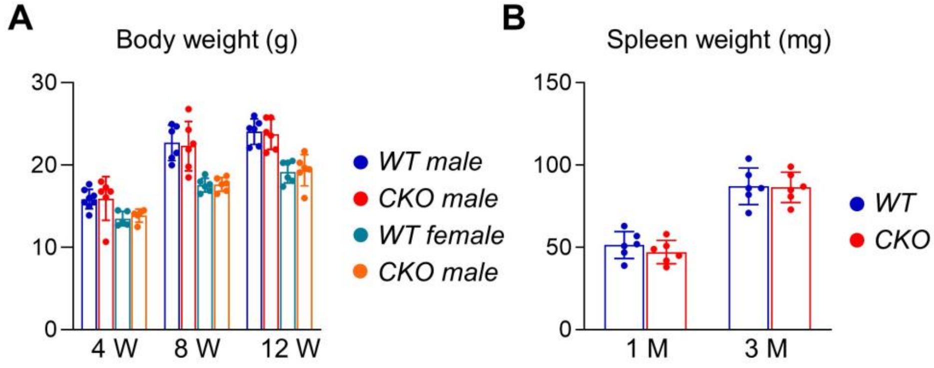


Figure S3. *RANKL CKO*$^{Adipoq}$ mice has normal body and spleen weight.

(A) Body weight was measured in male and female mice at 4, 8, and 12 weeks of age. n=5-8 mice/group.

(B) Spleen weight was measured at 1 and 3 months of age. n=6 mice/group.

**Figure S4**

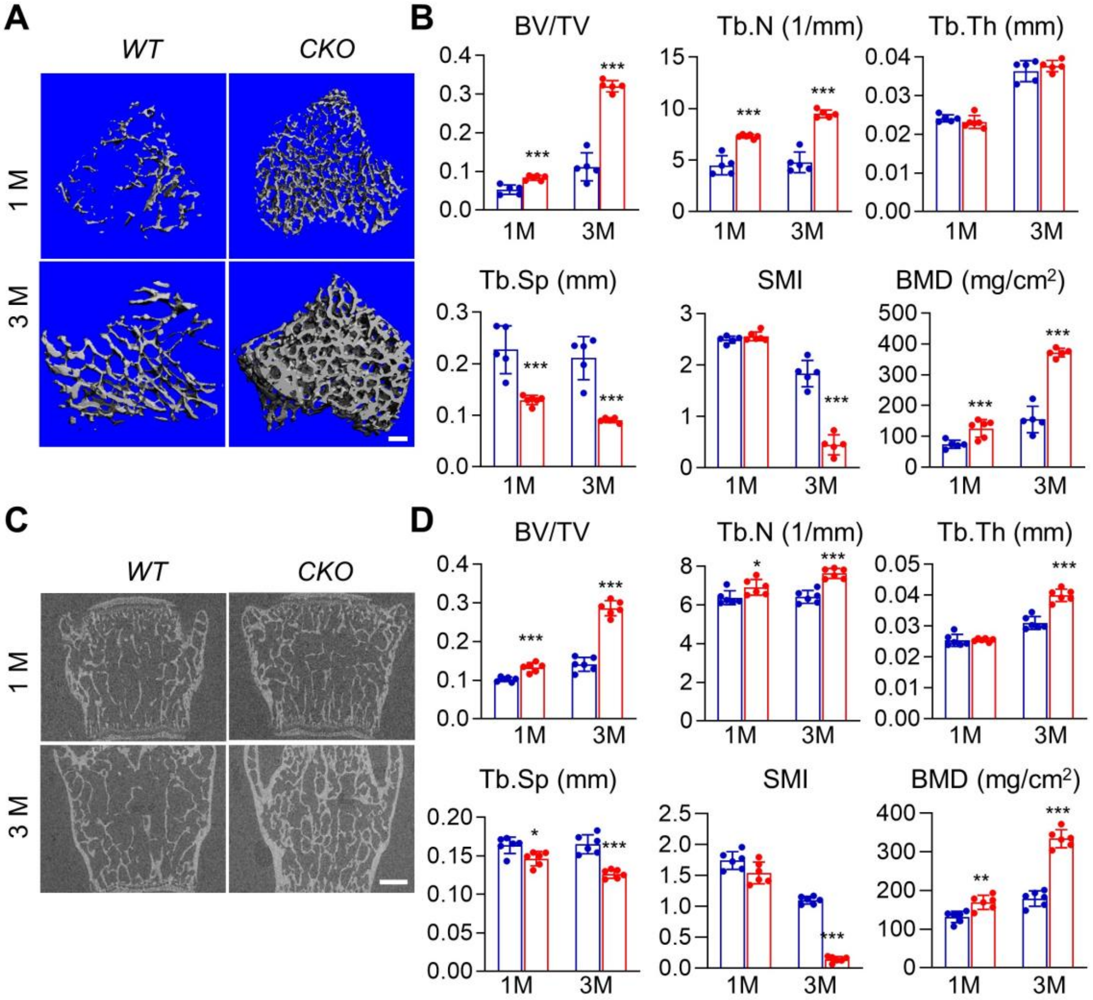


Figure S4. Female *RANKL CKO*$^{Adipoq}$ mice also display osteopetrosis phenotype.

(A) 3D microCT reconstruction of metaphyseal region in *WT* and *CKO* tibiae. Scale bar=200 μm.

(B) MicroCT measurement of trabecular bone structural parameters from the secondary spongiosa region. BV/TV: bone volume fraction; Tb.N: trabecular number; Tb.Th: trabecular thickness; Tb.Sp: trabecular separation; SMI: structural model index; BMD: bone mineral density. n=5-6 mice/group.

(C) 2D microCT reconstruction of *WT* and *CKO* vertebrates. Scale bar= 500μm.

(D) MicroCT measurement of trabecular bone structural parameters. n=5-6 mice/group.

*: $p<0.05$, **: $p<0.01$, ***: $p<0.001$ *CKO* vs *WT*.

**Figure S5**

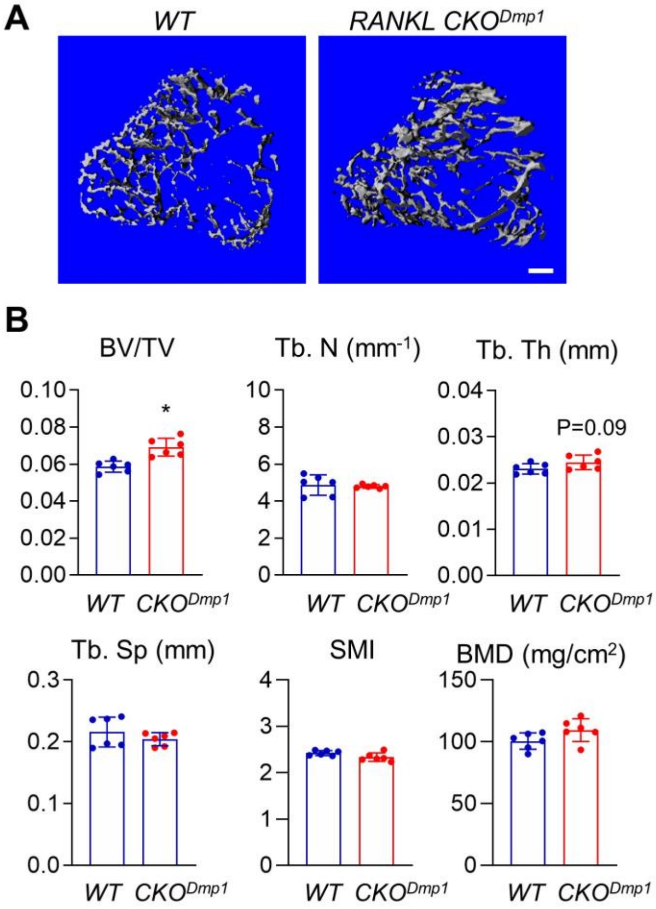


Figure S5. Mice with osteocyte-specific RANKL deficiency have a minor increase in trabecular bone mass at 1 month of age.

(A) 3D microCT reconstruction of metaphyseal region in *WT* and *RANKL CKO*$^{Dmp1}$ mouse tibiae. Scale bar=200 μm.

(B) MicroCT measurement of trabecular bone structural parameters from the secondary spongiosa region. BV/TV: bone volume fraction; Tb.N: trabecular number; Tb.Th: trabecular thickness; Tb.Sp: trabecular separation; SMI: structural model index; BMD: bone mineral density. *: $p<0.05$ *CKO* vs *WT*. n=6 mice/group.

**Figure S6**

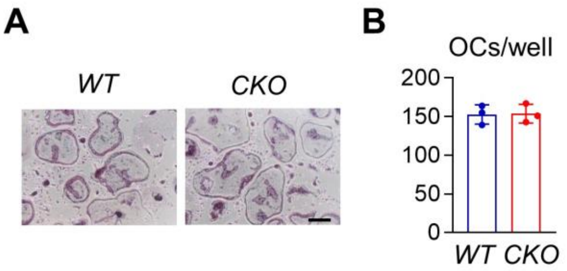


Figure S6. Osteoclast progenitors are normal in *RANKL CKO*$^{Adipoq}$ mice.

(A) Representative images of osteoclast culture derived from *WT* and *RANKL CKO*$^{Adipoq}$ BMCs at 7 days after addition of RANKL and M-Csf. Scale bar=200 μm.

(B) Quantification of TRAP$^{+}$ multinucleated cells per well. n=3 mice/group.

**Figure S7**

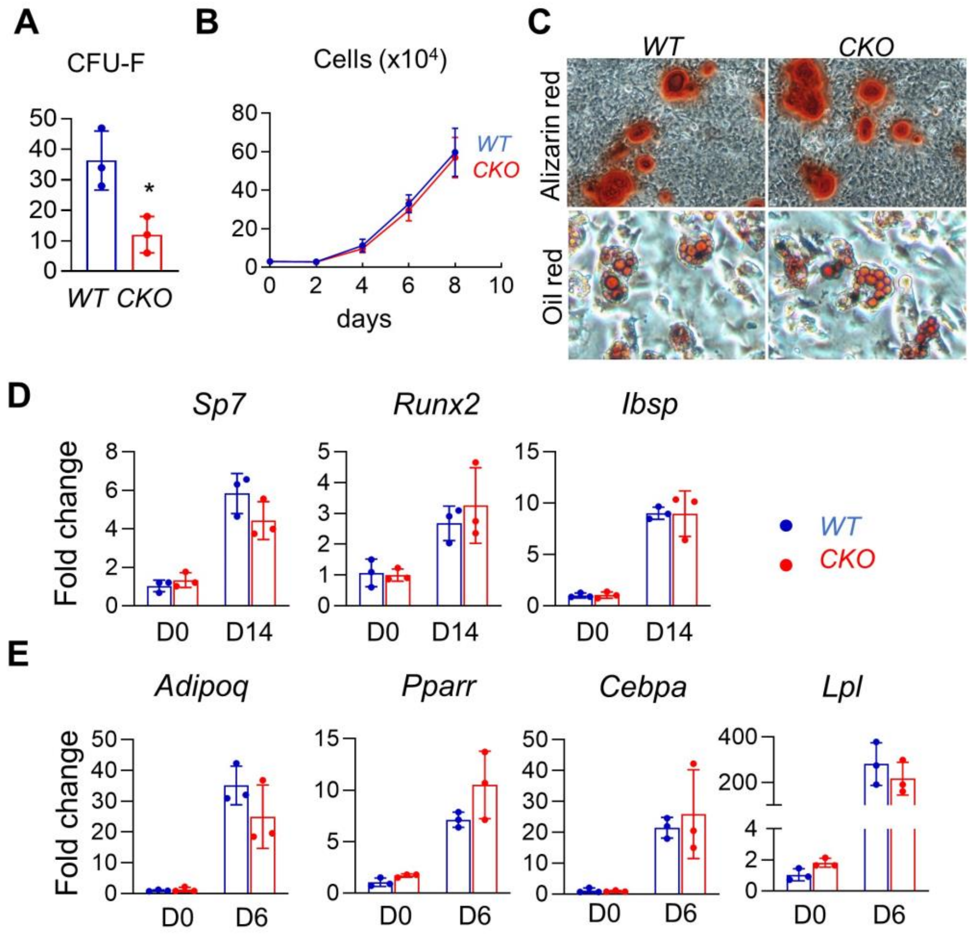


Figure S7. *RANKL CKO*$^{Adipoq}$ mice have a reduced pool of bone marrow mesenchymal progenitors.

(A) CFU-F assay of mesenchymal progenitors from femoral bone marrow of *WT* and *CKO* mice. *: $p<0.05$, *CKO* vs *WT*. n=3 mice/group.

(B) Growth curves of bone marrow mesenchymal progenitors in culture.

(C) Representative Alizarin red and Oil red staining of cells after cultured in osteogenic and adipogenic medium, respectively.

(D) qRT-PCR analysis of osteogenic marker gene expression in cells after 2 weeks of osteogenic differentiation.

(E) qRT-PCR analysis of adipogenic marker gene expression in cells after 1 weeks of adipogenic differentiation.

**Figure S8**

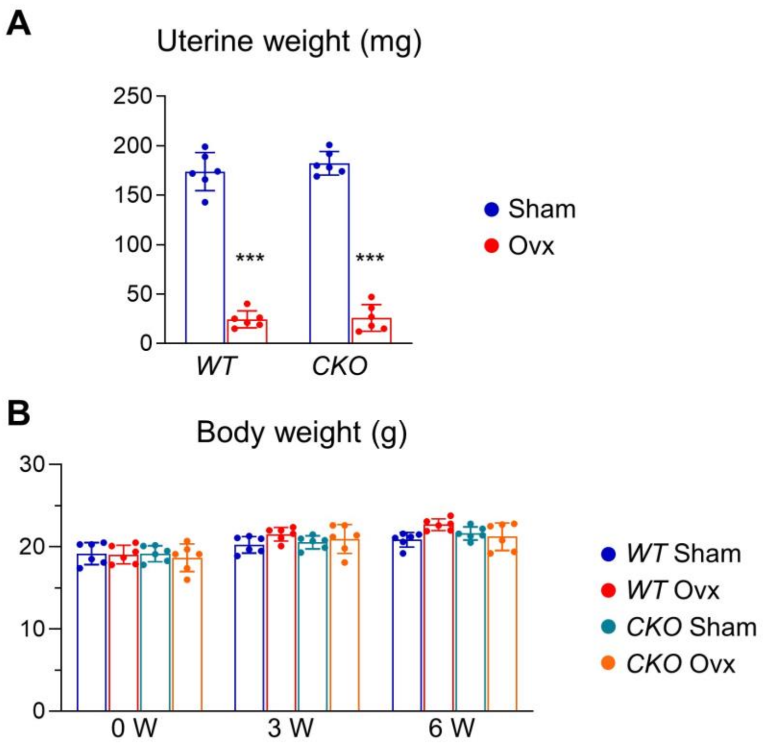


Figure S8. Examination of mouse uterine and body weight weight after ovx.

(A) Uterine weight of *WT* and *CKO* mice after ovx. ***: $p<0.001$ *CKO* vs *WT*. n=6 mice/group.

(B) Body weight of *WT* and *CKO* mice is recorded at different time points after ovx.

**Table S3.** Mouse real-time PCR primer sequences used in this study.

| Gene | Forward primer | Reverse primer |
|---|---|---|
| *β-actin* | 5'-TCCTCCTGAGCGCAAGTACTCT-3' | 5'-CGGACTCATCGTACTCCTGCTT-3' |
| *Tnfsf11* | 5'-GGAAGCGTACCTACAGACTA-3' | 5'-TGCTCCCTCCTTTCATCA-3' |
| *Sp7* | 5'-AGAGGTTCACTCGCTCTGACGA-3' | 5'-TTGCTCAAGTGGTCGCTTCTG-3' |
| *Runx2* | 5'-TAAAGTGACGGACGGTCCC-3' | 5'-TGCGCCCTAAATCACTGAGG-3' |
| *Ibsp* | 5'-ACCAGTTATGGCACCACGACA-3' | 5'-TCAACCGTGCTGCTCTTTCTG-3' |
| *Adipoq* | 5'-AAAGGAGAGCCTGGAGAA-3' | 5'-GAATGGGTACATTGGGAACA-3' |
| *Pparg* | 5'-CCAGCGTGAAGCCAGAGTAG-3' | 5'-ACCGTGGCTGTGCTCATCCT-3' |
| *Cebpa* | 5'-CAAGAACAGCAACGAGTACCG-3' | 5'-GTCACTGGTCAACTCCAGCAC |
| *Lpl* | 5'-GGGAGTTTGGCTCCAGAGTTT-3' | 5'-TGTGTCTTCAGGGGTCCTTAG |
